# Reduced Cdc14 phosphatase activity impairs septation, hyphal differentiation and pathogenesis and causes echinocandin hypersensitivity in *Candida albicans*

**DOI:** 10.1101/2022.09.29.510203

**Authors:** Kedric L. Milholland, Ahmed AbdelKhalek, Kortany M. Baker, Smriti Hoda, Andrew G. DeMarco, Noelle H. Naughton, Angela N. Koeberlein, Gabrielle R. Lorenz, Kartikan Anandasothy, Antonio Esperilla-Muñoz, Sanjeev K. Narayanan, Jaime Correa-Bordes, Scott D. Briggs, Mark C. Hall

**Author notes:** **Corresponding author – Email:**.

## Abstract

The Cdc14 phosphatase family is highly conserved in fungi. In *Saccharomyces cerevisiae,* Cdc14 is essential for down-regulation of cyclin-dependent kinase activity at mitotic exit. However, this essential function is not broadly conserved and requires a small fraction of normal Cdc14 activity. It remains unclear what fungal Cdc14 functions require high Cdc14 activity. We identified an invariant motif in the disordered C-terminal tail of fungal Cdc14 enzymes that is required for full enzyme activity. Mutation of this motif reduced Cdc14 catalytic rate and provided a tool for studying the biological significance of high Cdc14 activity. A *S. cerevisiae* strain expressing the reduced-activity hypomorphic mutant allele (*cdc14^hm^*) as the sole source of Cdc14 exhibited an unexpected sensitivity to cell wall stresses, including chitin-binding compounds and echinocandin antifungal drugs. Sensitivity to echinocandins was also observed in *Schizosaccharomyces pombe* and *Candida albicans* strains lacking *CDC14*, suggesting this phenotype reflects a conserved function of Cdc14 orthologs in mediating fungal cell wall integrity. In *C. albicans*, the orthologous *cdc14^hm^* allele was sufficient to elicit echinocandin hypersensitivity and perturb cell wall integrity signaling. It also caused striking abnormalities in septum structure and the same cell separation and hyphal differentiation defects previously observed with *cdc14* gene deletions. Since hyphal differentiation is important for *C. albicans* pathogenesis, we assessed the effect of reducing Cdc14 activity on virulence in *Galleria mellonella* and mouse models of invasive candidiasis. Partial reduction in Cdc14 activity via *cdc14^hm^* mutation severely impaired *C. albicans* virulence in both assays. Our results reveal that high Cdc14 activity promotes fungal cell wall integrity and, in *C. albicans*, is needed to orchestrate septation and hyphal differentiation, and for pathogenesis. Cdc14 may therefore be worth future exploration as an antifungal drug target.

**AUTHOR SUMMARY:** Invasive fungal infections are a serious concern for the immune-compromised. Antifungal drugs to treat invasive infections are limited and pathogens are developing resistance to them. Novel targets for antifungal drug development are needed. In this study we developed a system to test if partial therapeutic reduction in activity of a protein phosphatase called Cdc14 could reduce virulence of the opportunistic human pathogen *Candida albicans.* This idea arose from prior studies in fungal pathogens of plants, where Cdc14 was unexpectedly required for host infection through an unknown mechanism. We found that successful *C. albicans* infections in two animal models of invasive candidiasis were dependent on high Cdc14 activity. Moreover, we made the surprising observation that integrity of the *C. albicans* cell wall is also dependent on high Cdc14 activity, with Cdc14-deficient cells becoming hypersensitive to cell wall-targeted antifungal drugs. We conclude that even modest reduction in Cdc14 activity could have therapeutic benefit for human fungal infections and possibly help overcome resistance to some antifungal drugs. Cdc14 structure and specificity are unique among phosphatases and highly conserved in pathogenic fungi, suggesting that highly selective inhibitors can be developed that would be useful against a broad range of fungal pathogens.

## INTRODUCTION

*CDC14* was originally identified in *Saccharomyces cerevisiae* as an essential gene required for the final stage of nuclear division (1). The Cdc14 protein was subsequently characterized as a protein phosphatase that fulfilled its essential function by terminating, and reversing the effects of, mitotic cyclin-dependent kinase (Cdk) activity to trigger mitotic exit (2). *CDC14* is widely conserved in the major eukaryotic lineages, with the exception of angiosperm plants (3–5), and Cdc14 catalytic domain structure, activity and specificity are also broadly conserved (4,6–8). For example, human CDC14 orthologs can complement *cdc14* mutant phenotypes in *S. cerevisiae* and *Schizosaccharomyces pombe* (9,10).

Cdc14 is related to the dual-specificity phosphatase (DSP) subfamily of protein tyrosine phosphatases (PTP), sharing the invariant HCX5R catalytic motif and two-step catalytic mechanism involving a covalent phospho-enzyme intermediate (11,12). Despite its clear evolutionary relationship to the DSP and PTP families, Cdc14 evolved a very strong preference for phosphoserine sites followed by the minimal sequence motif P-X-K (P-X-R to a lesser extent) in an ancestral eukaryote, making it functionally a phosphoserine phosphatase. This unique specificity for a PTP family member arose from an apparent duplication of the DSP domain, which created a novel substrate binding groove along a domain interface adjacent to the active site (4,6,7). The tandem arrangement of DSP folds, and the specificity it imparts, are signature features of Cdc14 enzymes across eukaryotes. The broad conservation implies Cdc14 specificity must fill an essential niche in cellular regulation of most eukaryotic species that cannot be provided by other prominent protein phosphatases. One hypothesis, based on work in the oomycete *Phytopthora infestans,* is that Cdc14 co-evolved with eukaryotic flagella (5), and subsequent studies have reported roles for Cdc14 enzymes in regulating aspects of cilia structure and function in vertebrates, including human cells (13–15). The fungal subkingdom Dikarya, which includes most fungal pathogens of plants and animals, represents a striking exception to the Cdc14-flagella correlation, as they have strictly retained *CDC14* during evolution despite having lost flagella.

Within Dikarya, Cdc14 functional characterizations have been conducted in several species, including the model yeasts *S. cerevisiae* and *S. pombe.* Interestingly, the originally described essential function in terminating Cdk activity and triggering mitotic exit in *S. cerevisiae* is not widely conserved, even in other ascomycetes. In fact, *CDC14* deletions in other fungal species often have modest impacts on vegetative cell division (16–21). Moreover, we and others found that the essential function in *S. cerevisiae* requires only a small fraction of natural Cdc14 expression (22–24). Cdc14 is an abundant phosphatase in *S. cerevisiae,* with protein levels comparable to subunits of *PP2A* phosphatases (25). The fact that a small fraction of expressed Cdc14 is sufficient for the essential function in *S. cerevisiae* implies that the majority of Cdc14 is necessary for other important cellular functions.

Cdc14 has been linked to regulation of diverse biological processes in model fungi, including centrosome duplication, mitotic spindle dynamics, chromosome segregation, DNA replication and repair, polarized growth, autophagy, and response to osmotic and genotoxic stresses (26). The conservation of many of these functions remains uncertain but it is now becoming clear that a highly conserved function of fungal Cdc14 phosphatases is regulation of cytokinesis and septation at the end of the cell cycle. The *S. pombe* Cdc14 ortholog, Clp1, does not regulate mitotic exit but plays multiple roles ensuring correct cytokinesis and septation (20,27). Blocking Cdc14 nuclear export in *S. cerevisiae* allowed normal mitotic exit regulation but resulted in cytokinesis defects (28). In the phytopathogenic fungi *Fusarium graminearum* and *Magnaporthe oryzae, CDC14* deletion impaired septum formation and the coordination between nuclear division and cytokinesis, leading to multi-nucleated conidial and hyphal compartments (16,17). Similar septation defects were observed after *CDC14* deletion in another plant pathogen, *Aspergillus flavus* (18). Finally, in the opportunistic human pathogen *Candida albicans, CDC14* deletion caused defective cell separation after cytokinesis and septation, resulting in connected chains of cells from failure to activate the Ace2 transcription factor (29). A number of Cdc14 substrates have been identified and implicated in cytokinesis and septation regulation in *S. cerevisiae* and *S. pombe* (30) but the relevant substrates remain unknown in most other fungal species.

Importantly, in studies of *CDC14* deletion-associated phenotypes in phytopathogenic fungi, severe reductions in virulence were observed, even when vegetative growth was only marginally impacted (16–18). The molecular mechanisms by which Cdc14 promotes plant infection are undefined. It is unknown if *CDC14* is also important for virulence in animal fungal pathogens, although the *CDC14* deletion in *C. albicans* impaired hyphal development, which is believed to be important for virulence (31,32). Collectively, these reports suggest that one or more conserved Cdc14 functions in fungi is generally important for host infection by pathogens.

In an attempt to better understand fungal Cdc14 functions unrelated to mitotic exit that require high level expression, we sought a way to partially reduce Cdc14 activity *in vivo.* We focused on the poorly conserved, and seemingly disordered, C-terminal tail, which is dispensable for phosphatase activity and the essential mitotic exit function in *S. cerevisiae* (11). In some Cdc14 orthologs this region has regulatory phosphorylation sites and elements that control cellular localization (28,33,34), but other specific functions have not been described. Of note, the isolated ScCdc14 catalytic domain lacking the C-terminal region exhibited reduced activity towards the generic phosphatase substrate *para*-nitrophenyl phosphate (pNPP) in an early characterization of Cdc14 enzyme activity (11). We subsequently observed a similar phenomenon with truncated Cdc14 enzymes from *F. graminearum* (16) and other fungal species (our unpublished data), suggesting that the C-terminal region does make a conserved contribution to full catalytic activity. Since the existing crystal structures of Cdc14 phosphatases lack the C-terminal tail (6,7), it is unclear how this might occur, but it suggests that modulation of the C-terminus might allow creation of reduced activity hypomorphic alleles that are otherwise normal and could be used to study the functional importance of high level Cdc14 expression.

In this report, we identify a short, invariant motif in the disordered C-terminus of fungal Cdc14 enzymes that stimulates Cdc14 activity and is a likely target for *in vivo* Cdc14 regulation. We characterized mutations in this motif in *S. cerevisiae* and *C. albicans.* The results revealed an unexpected new role for Cdc14 in promoting fungal cell wall integrity and demonstrated that high level Cdc14 expression is critically important for *C. albicans* septation, hyphal development, and pathogenesis.

## RESULTS

### A conserved motif in the fungal Cdc14 C-terminus is required for full catalytic activity

The observation that the isolated catalytic domains of multiple fungal Cdc14 orthologs have reduced enzyme activity raised the possibility of creating hypomorphic mutants to test the importance of full Cdc14 activity. Using *S. cerevisiae* Cdc14 (ScCdc14), we set out to identify regions of the ~200 amino acid C-terminal tail that specifically contribute to enzyme activity. We purified full length ScCdc14(1-551), the catalytic domain truncation ScCdc14(1-374) and a truncation of intermediate length, ScCdc14(1-449) (**Fig 1A and Fig S1A**) and compared activity under steady-state conditions. Consistent with prior reports (11,16), ScCdc14(1-374) reaction rate towards pNPP was greatly reduced compared to ScCdc14(1-551) (**Fig 1B**). ScCdc14(1-449) activity was nearly identical to ScCdc14(1-551), demonstrating that the sequence between amino acids 374 and 449 is required for full activity. We used ScCdc14(1-449) to represent full, or “wild-type”, activity in subsequent experiments. The same analysis with a biologically-relevant phosphopeptide substrate derived from a known ScCdc14 substrate site in Yen1 (36) revealed a similar large reduction in *k_cat_* for ScCdc14(1-374) compared to ScCdc14(1-449), indicating that the difference in activity is not substrate-dependent (**Fig 1C**).

**Figure 1.**
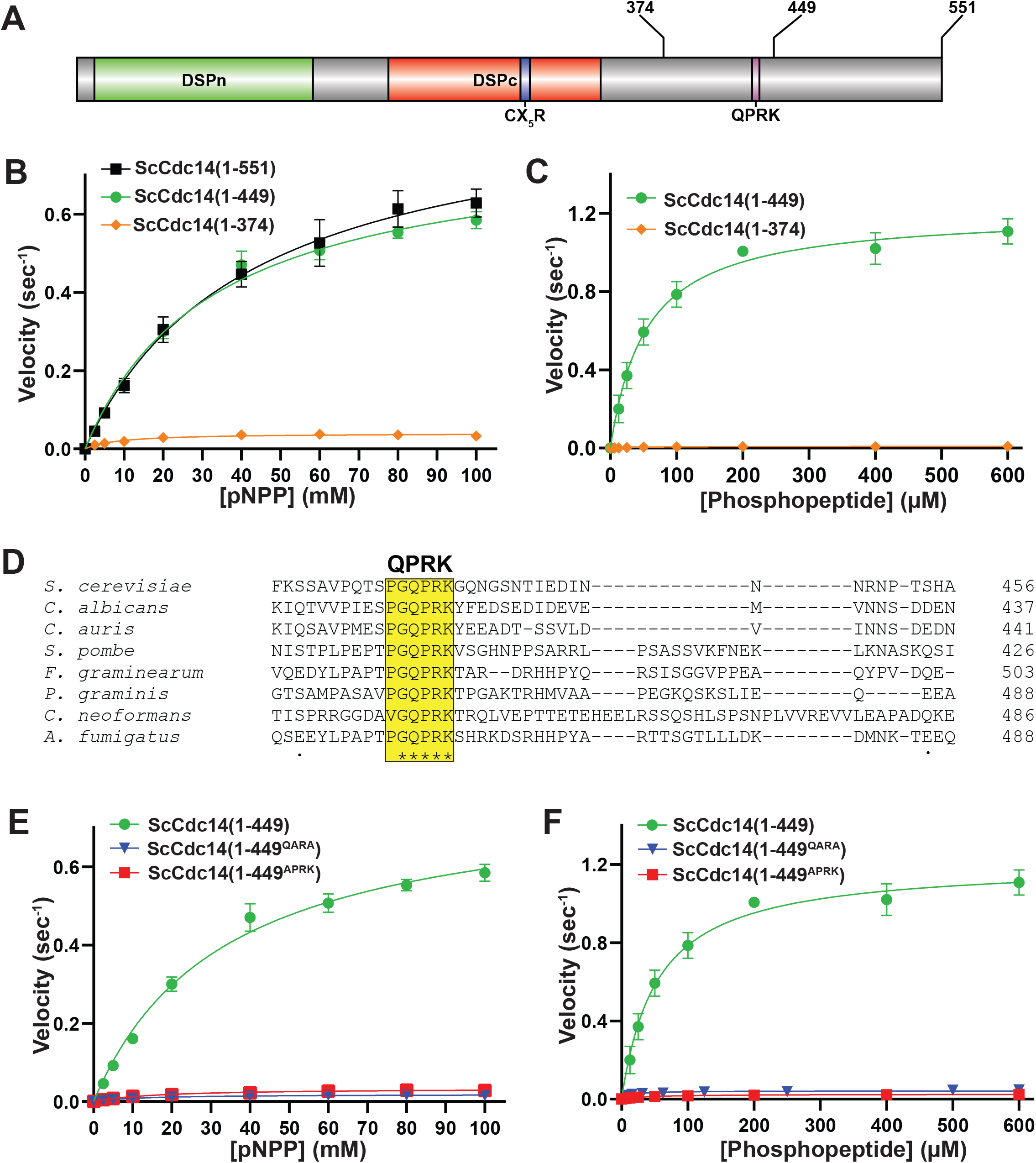
A conserved motif in Cdc14’s disordered C-terminal tail is required for full catalytic activity. (**A**) Domain map of Cdc14. DSPn and DSPc are the N- and C-terminal dual-specificity phosphatase domains, respectively. CX5R is the invariant PTP family catalytic motif. Sites of truncations and the invariant QPRK motif are indicated in the disordered C-terminus following DSPc. (**B**) Steady-state kinetic analyses with full length ScCdc14(1-551) and the indicated truncation variants towards pNPP yielded kcat values of 0.91 (1-551), 0.80 (1-449) and 0.04 (1-374) sec^-1^. (**C**) Same as (B) with phosphopeptide substrate HT[pS]PIKSIG, yielding *k_cat_* values of 0.64 (1-449) and 0.01 (1-374) sec^-1^. (**D**) Multiple sequence alignment of fungal Cdc14 orthologs from the indicated species was generated by Clustal Omega using default settings. Only a portion of the C-terminal region showing the invariant QPRK motif is shown. (**E**) Steady-state kinetic analyses comparing wild-type ScCdc14 and QPRK motif mutants towards pNPP; *k_cat_* values were 0.02 (1-449^QARA^) and 0.03 (1-449^APRK^) sec^-1^. (**F**) Same as (E) with the phosphopeptide substrate; *k_cat_* values were 0.02 (1-449^QARA^) and 0.03 (1-449^APRK^) sec^-1^. The ScCdc14(1-449) data in panels E and F are identical to panel C. All kinetic data in panels B, C, E, and F are averages of at least 3 independent trials with standard deviation error bars. Fit lines and *k_cat_* values were generated in GraphPad Prism using the Michaelis-Menten function.

A multiple sequence alignment of diverse fungal Cdc14 orthologs revealed a single, short invariant sequence within the otherwise poorly conserved C-terminus (**Fig 1D**) that was not observed in animal Cdc14 sequences (**Fig S2**). Part of this sequence, QPRK, strongly resembles the optimal substrate recognition motif of Cdc14 enzymes, SPxK (“x” is preferentially a basic amino acid) (7,8,36), suggesting it could influence catalysis by interacting with the active site as a pseudosubstrate. Furthermore, this motif is within residues 374-449 required for full activity. We substituted Ala either for the Pro and Lys residues, analogous to two critical positions for Cdc14 substrate recognition (7,8,36), or for the Gln, since this residue would occupy the pSer binding pocket if the motif were acting as a pseudosubstrate. The resulting ScCdc14(1-449^QARA^) and ScCdc14(1-449^APRK^) variants exhibited *k_cat_* decreases similar to ScCdc14(1-374) with both pNPP (**Fig 1E**) and phosphopeptide (**Fig 1F**) substrates. Mutation of QPRK did not affect substrate specificity as activity was still dependent on the same major substrate features as wild-type Cdc14 enzymes (**Fig S1B**). We conclude the QPRK motif makes a significant contribution to enzyme turnover rate and is responsible for the observed rate difference between catalytic domain truncations and full-length enzymes.

### QPRK motif mutation causes sensitivity to cell wall stress in *S. cerevisiae*

The strict conservation of the QPRK motif across the fungal kingdom suggests it is biologically important. To characterize its biological significance, we generated a haploid*S. cerevisiae* strain expressing the **<u>h</u>**ypo**<u>m</u>**orphic *cdc14^QARA^* allele (hereafter called *cdc14^hm^)* as the sole source of Cdc14 from the natural *CDC14* locus using the *delitto perfetto* approach (37). In liquid culture, the mutant strain grew at a similar rate to the wild-type parental strain at low cell density but slowed somewhat as cell density increased and ultimately terminated at a slightly lower optical density (**Fig S3A**). Thus, the essential mitotic exit function was not disrupted by the reduction in Cdc14 catalytic rate, consistent with our prior observations that it requires a small fraction of normal Cdc14 activity (22). We performed agar plate spotting assays in the presence of a variety of cellular stress conditions to determine if reducing Cdc14 activity sensitizes cells to stress. *cdc14^hm^* colony size was slightly smaller than wild-type on untreated control plates but otherwise grew similarly and *cdc14^hm^* was no more sensitive than wild-type to genotoxic, oxidative, and osmotic stresses (**Fig 2A** and **Fig S3B-D**). The temperature-sensitive *cdc14-3* allele was previously shown to impart some sensitivity to calcofluor white (CFW), a chitin-binding compound that causes cell wall stress (38). We confirmed this observation and found that *cdc14^hm^* exhibited strong sensitivity to CFW (**Fig 2B**) and another chitin-binding compound, Congo red (**Fig S3E**). To determine if this sensitivity was unique to chitin binding chemicals we performed spotting assays in the presence of echinocandin drugs, which cause cell wall stress by inhibiting synthesis of the major cell wall polymer (1,3)-β-d-glucan (39). *cdc14^hm^* was acutely sensitive to both micafungin and caspofungin compared to wild-type (**Fig 2C-D**). *cdc14^hm^* was also sensitive to growth at high temperature, another condition that generates cell wall stress (40) (**Fig S3F**). Cell wall stress sensitivity was observed when *cdc14^hm^,* but not *CDC14,* was expressed from a CEN plasmid in a *cdc14Δ* background **(Fig S3G).**Sensitivity in *cdc14^hm^* was complemented by transformation with a CEN plasmid expressing wild-type *CDC14* **(Fig S3H),** suggesting it is a recessive loss of function phenotype caused by the reduced activity. Micafungin sensitivity was rescued in the presence of sorbitol, an osmotic stabilizer known to suppress cell wall integrity defects (41), suggesting that reduced Cdc14 activity compromises cell wall integrity (**Fig 2E**).

**Figure 2.**
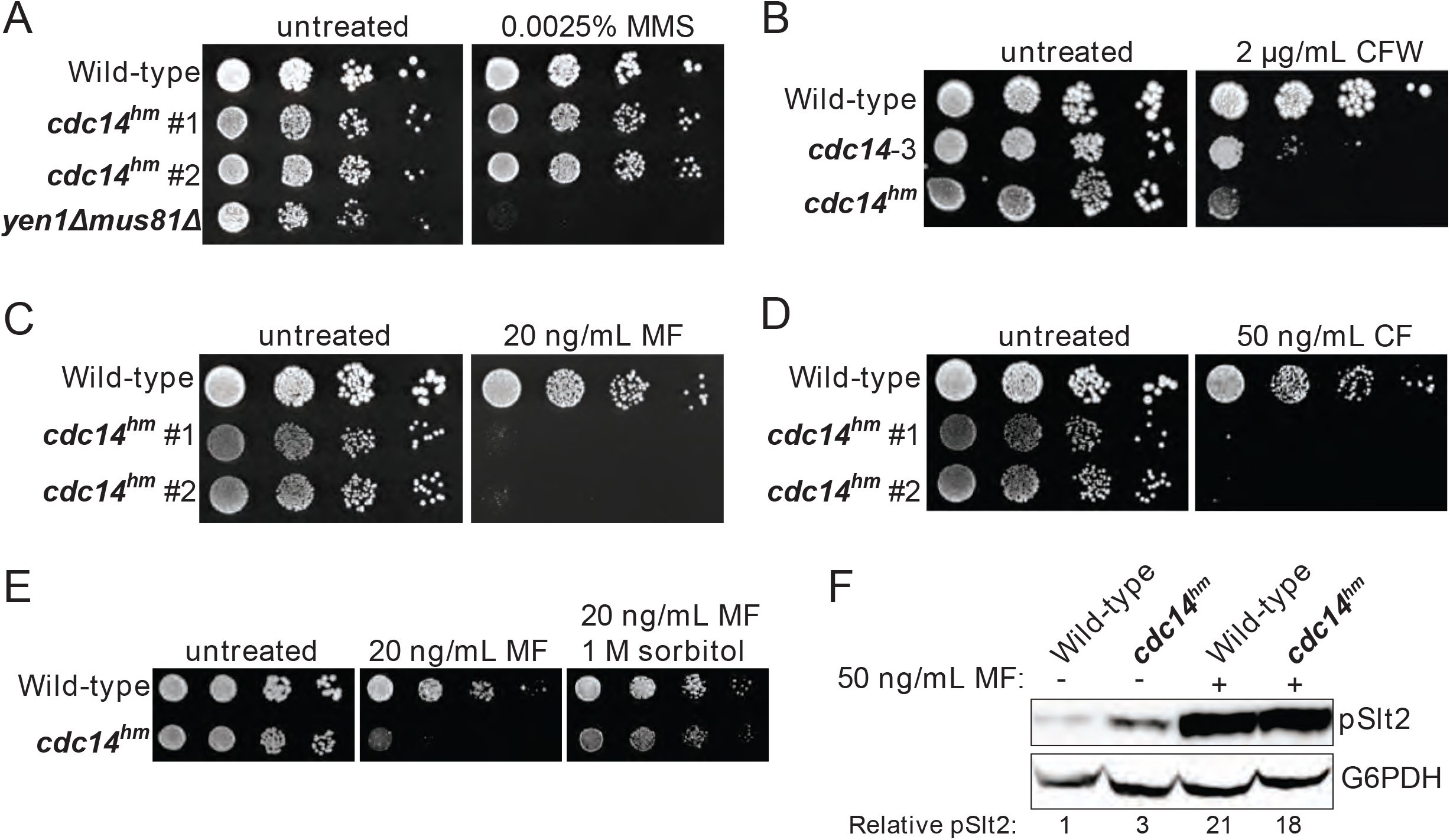
QPRK mutation causes sensitivity to cell wall stress in *S. cerevisiae*. (**A-E**) Cultures of the indicated *S. cerevisiae* strains were serially diluted and spotted on YPAD agar plates containing indicated concentrations of (**A**) methyl methanesulfonate (MMS), (**B**) calcofluor white (CFW), (**C**) micafungin (MF), (**D**) caspofungin (CF), and (**E**) MF or MF + sorbitol. “#1” and “#2” refer to independent isolates of *cdc14^hm^. yen1Δ mus81Δ* is a positive control for MMS sensitivity (87). All plates were incubated at 30°C for 72 hours. Images are representative of three biological replicates. “Wild-type” is W303. (**F**) Immunoblot detecting phosphorylated, active form of Slt2 (pSlt2) from cell lysates of mid-log phase cells with anti-p44/42 MAP kinase antibody. pSlt2 signals were corrected based on the G6PDH (glucose-6-phosphate dehydrogenase) load control and then normalized to the untreated wild-type sample to generate the Relative pSlt2 value, which is an average of two independent trials.

Cell wall damage activates the cell wall integrity (CWI) signaling pathway (42). Activation of the CWI pathway can be monitored by immunoblotting for the phosphorylated CWI MAP kinase, Slt2. We compared phospho-Slt2 levels in log phase wild-type and *cdc14^hm^* cultures in the absence and presence of micafungin. *cdc14^hm^* showed a 3-fold increase in phospho-Slt2 in the absence of micafungin, but both wild-type and *cdc14^hm^* strains showed similar Slt2 activation after brief treatment with micafungin (**Fig 2F)**. We also monitored expression of *FKS2,* which encodes a catalytic subunit of (1,3)-β-d-glucan synthase that is induced by CWI signaling during cell wall stress (43–45). Consistent with the pSlt2 result, *FKS2* transcript level was 2-fold higher in *cdc14^hm^* compared to the isogenic wild-type strain in the absence of micafungin (**Fig S3I**). These results are consistent with a cell wall integrity defect, but mostly normal CWI signaling, in *cdc14^hm^ S. cerevisiae* cells.

### Cdc14-dependent cell wall stress hypersensitivity is conserved in *S. pombe* and *C. albicans*

To test if Cdc14-dependent sensitivity to cell wall stress is conserved in other fungal species, we performed spotting assays with *CDC14* deletion strains of *S. pombe* and *C. albicans. S. pombe clp1Δ* exhibited clear sensitivity to caspofungin, but not calcofluor white, presumably due to the lack of chitin in *S. pombe* cell walls (46) **(Fig 3A**). *C. albicans cdc14Δ/Δ* exhibited strong sensitivity to calcofluor white, caspofungin, and micafungin (**Fig 3B**). These observations suggest that Cdc14 makes a broadly conserved contribution to fungal cell wall integrity.

**Figure 3.**
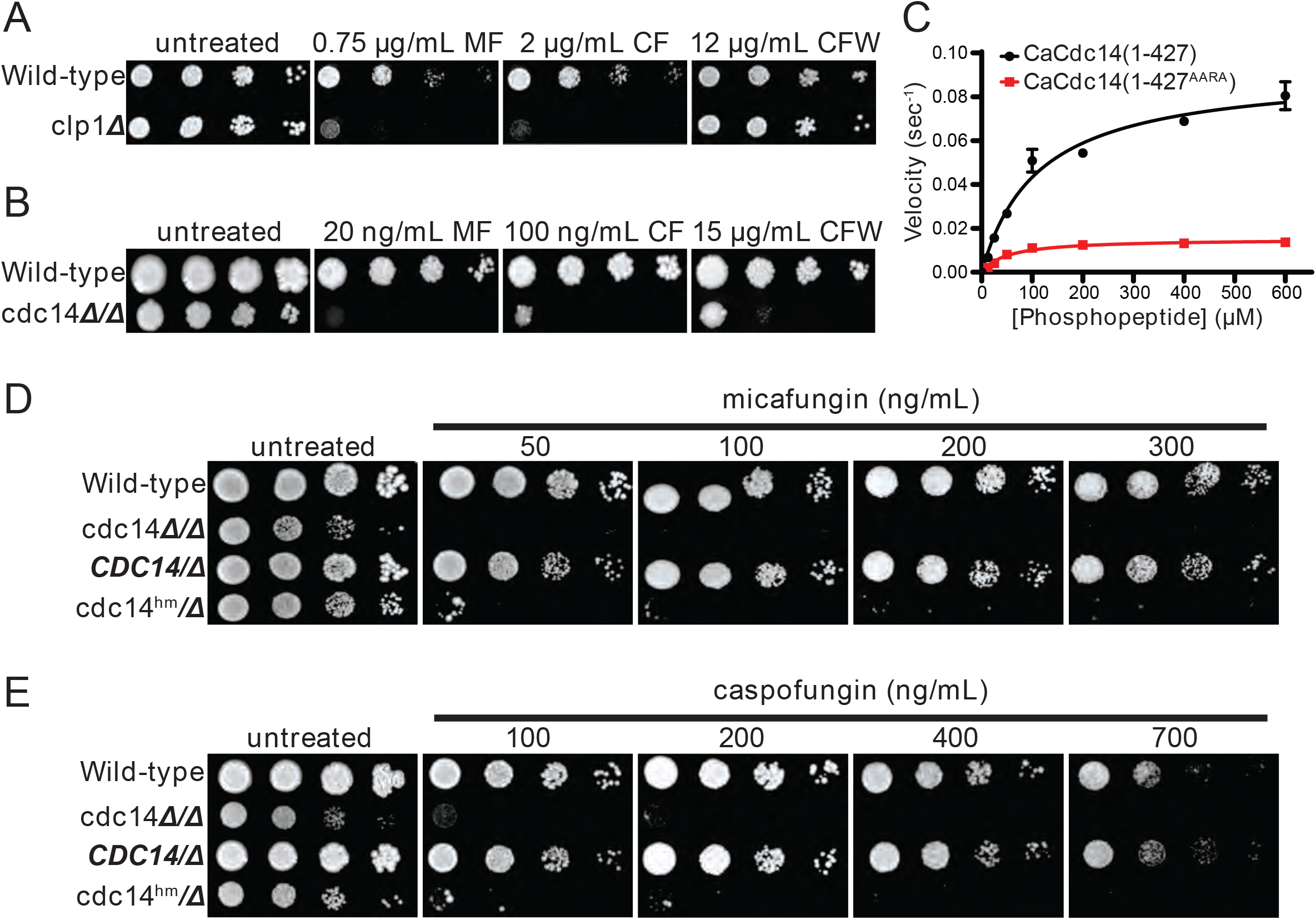
Cdc14-dependent cell wall stress sensitivity is conserved in other fungal species. Liquid cultures of the indicated *S. pombe* (**A**) and *C. albicans* (**B**) strains were serially diluted and spotted on YES or YPD agar plates, respectively, containing the indicated concentrations of micafungin (MF), caspofungin (CF), or calcofluor white (CFW). *CLP1* is the *S. pombe* homolog of *CDC14.* (**C**) Steady-state kinetic analysis of CaCdc14(1-427) and CaCdc14(1-427^AARA^) using the Yen1 phosphopeptide substrate. *k_cat_* values were 0.092 (1-427) and 0.015 (1-427^AARA^) sec^-1^. Data are means from three independent trials and error bars are standard deviations. Liquid cultures of the indicated *C. albicans* strains were serially diluted and spotted on YPD agar plates containing the indicated concentrations of micafungin (**D**) or caspofungin (**E**). Plates were incubated at 30 °C for 3-5 days (until mature colonies were visible in the wild-type strain). Images are representative of three independent trials.

We next tested if the QPRK motif functions similarly in *C. albicans* Cdc14 (CaCdc14) and if mutating it is sufficient to cause cell wall stress sensitivity, as in *S. cerevisiae.* To our knowledge, CaCdc14 has not been previously purified and characterized. We purified recombinant truncated CaCdc14 (residues 1-427, which retains the QPRK motif), as full-length protein was insoluble, and created a variant with Ala substitutions at the Gln, Pro, and Lys positions of the QPRK motif (**Fig S4A**). CaCdc14(1-427) exhibited the same major substrate preferences as ScCdc14 and other Cdc14 orthologs (**Fig S4B**), consistent with its identical active site composition (**Fig S2**). Steady-state kinetic analyses with the Yen1 phosphopeptide substrate revealed a 6-fold reduction in *k_cat_* for CaCdc14(1-427^AARA^) compared to CaCdc14(1-427) (**Fig 3C**). Thus, the QPRK motif makes a similar, albeit somewhat weaker, contribution to CaCdc14 phosphatase activity.

To study the biological significance of the QPRK motif in *C. albicans* we complemented *cdc14Δ/Δ* with either *cdc14^hm^-3xHA* (where *cdc14^hm^* encodes the QPRK➔AARA substitutions) or *CDC14-3xHA* (**Fig S4C)***. cdc14^hm^* growth rate was intermediate between the wild-type and *cdc14Δ/Δ* strains, consistent with a partial loss of activity and function (**Fig S4D**). In agar plate spotting assays *cdc14^hm^/Δ* displayed hypersensitivity to a range of echinocandin concentrations, similar to *cdcΔ4Δ/Δ*, whereas *CDC14/Δ* behaved like the *CDC14/CDC14* wild-type parent strain (**Fig 3D-E**). Thus, echinocandin hypersensitivity requires only a partial reduction in Cdc14 activity. As in *S. cerevisiae,* hypersensitivity was specific to cell wall stress as *cdc14Δ/Δ* and *cdc14^hm^/Δ* were mostly insensitive to genotoxic, oxidative, and osmotic stress (**Fig S4E**).

### Reduced Cdc14 activity causes elevated CWI signaling and alters cell wall stress-induced gene expression in *C. albicans*

We next tested if CWI signaling was altered in Cdc14-deficient *C. albicans* strains, similar to *S. cerevisiae.* In *C. albicans,* the anti-p44/42 MAPK antibody used to detect the activated form of the CWI MAP kinase produced several bands in our hands. We used the GRACE collection strains (47) for *MKC1* (the *C. albicans* ortholog of *S. cerevisiae* Slt2) and the closely related MAP kinase *CEK1* to ensure we were monitoring Mkc1 activation (**Fig S5**). In the absence of added stress, liquid cultures of both *cdc14Δ/Δ* and *cdc14^hm^/Δ* displayed an equal elevation in activating phosphorylation of Mkc1 compared to *CDC14/CDC14* (**Fig 4A-B**). Interestingly, *CDC14/Δ* cells displayed an intermediate elevation in phospho–Mkc1 signal, suggesting that even *CDC14* haplo-insufficiency can cause some chronic cell wall abnormality. The reason for this is unclear, but it suggests that CWI is particularly sensitive to Cdc14 activity level in *C. albicans*. In contrast to the basal state, when cultures were briefly challenged with micafungin, Mkc1 activation was significantly lower in both *cdc14Δ/Δ* and *cdc14^hm^/Δ* compared to *CDC14/CDC14* and *CDC14/Δ* (**Fig 4A-B**). This suggests that Cdc14 activity may also play a role in normal CWI pathway activation in response to certain cell wall stress signals.

**Figure 4.**
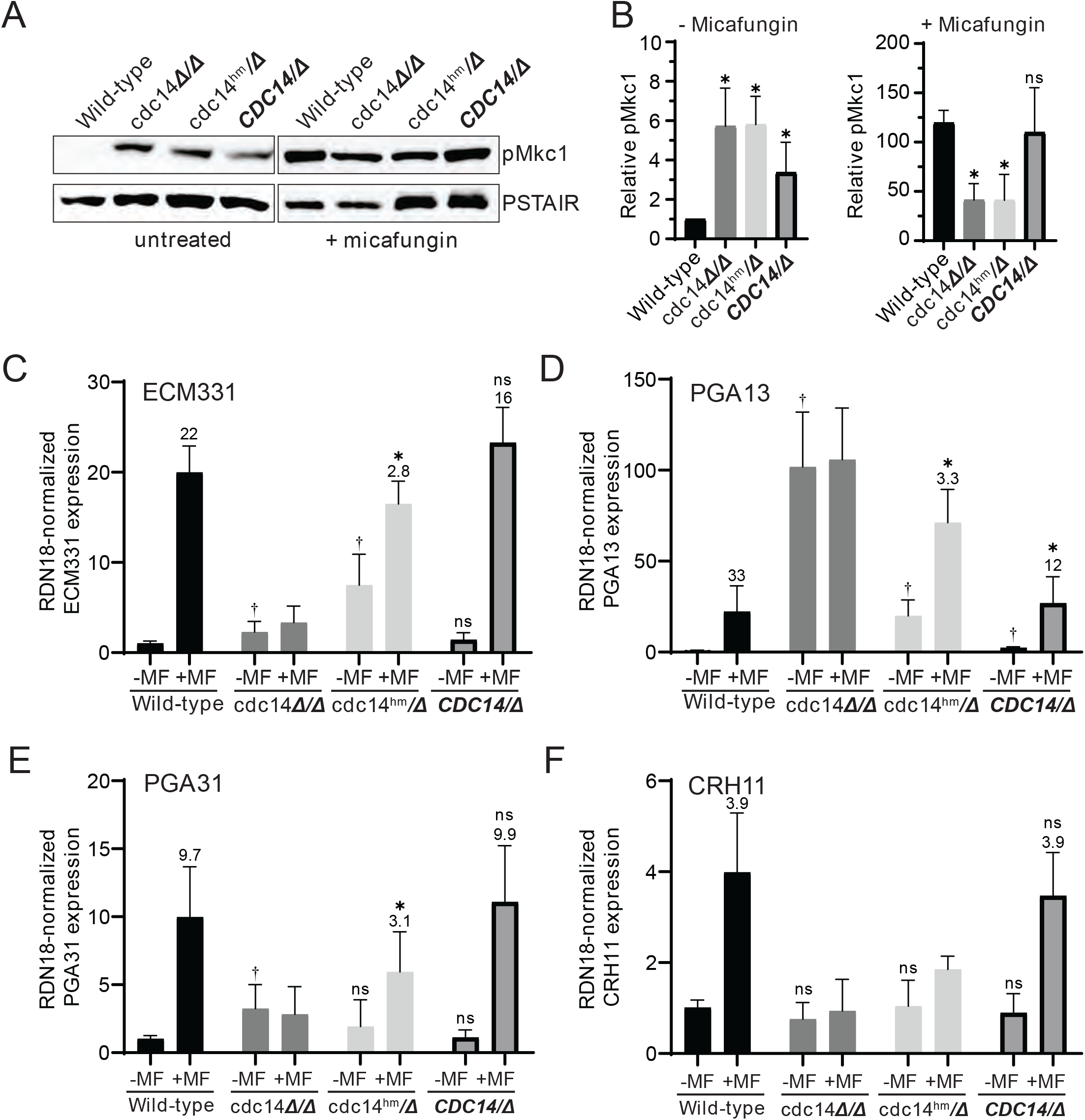
Reduced Cdc14 activity causes elevated CWI signaling and perturbs cell wall gene expression in *C. albicans*. (**A**) Phosphorylation of Mkc1 (pMkc1) was detected in whole cell extracts from mid-log phase cultures of the indicated *C. albicans* strains by anti-p44/p42 MAP kinase immunoblotting. Saturated cultures were back-diluted to OD_600_ = 0.01 and grown to mid-log phase. A portion of the untreated culture was collected and the remainder was treated with 50 ng/mL micafungin (MF) for 30 min. anti-PSTAIR is a loading control. (**B**) pMkc1 chemiluminescence signals from (A) were normalized to PSTAIR and plotted as a fold-increase above the untreated wild-type signal. Values are means from three independent experiments and error bars are standard deviations. Unpaired T-tests assuming equal variance comparing pMkc1 level in each strain to wild-type were used to determine statistical significance - * = *p* < 0.05; ns = not significant (*p* > 0.05). (**C-F**) Relative mRNA levels of selected cell wall stress-responsive genes in mid-log phase cultures of *C. albicans* strains either untreated (-) or treated (+) with 50 ng/mL MF for 1 h were measured by qRT-PCR. Transcript levels were normalized to *RDN18* and set relative to the untreated wild-type sample. Data are averages of at least 5 biological trials and error bars are standard deviations. Outliers were identified by Grubbs test and removed if detected. Unpaired T-tests assuming equal variance were used to compare 1) the basal (untreated) expression in each strain with wild-type († =*p* < 0.05) and 2) untreated with MF-treated samples of each strain. If a significant increase was observed with MF treatment, then the average fold-increase was added above the bar and an additional T-test was performed to determine if fold-increase was significantly different than wild-type fold-increase (* =*p* < 0.05). ns = not significant (*p* > 0.05).

To test if Cdc14 activity affects gene expression downstream of Mkc1 we used qRT-PCR to measure transcript levels of several cell wall stress-responsive genes (48) in the absence and presence of micafungin. Consistent with the phospho-Mkc1 immunoblotting results, *ECM331, PGA13,* and *PGA31* transcripts were significantly elevated in *cdc14Δ/Δ* and/or *cdc14^hm^/Δ* strains compared to *CDC14/CDC14* and *CDC14/Δ* in the absence of micafungin, although *CRH11* showed no change (**Fig. 4C-F**). Interestingly, after brief micafungin treatment *cdc14Δ/Δ* cells failed to induce expression of any of the four genes and the fold-induction in *cdc14^hm^/Δ* was significantly reduced compared to wild-type strains. This effect was not due to the genes already being maximally induced since three out of four transcript levels were lower in micafungin-treated *cdc14Δ/Δ* and *cdc14^hm^/Δ* compared to *CDC14/CDC14* and *CDC14/Δ*. Thus, Cdc14 activity is important for expression of at least some cell wall stress-responsive genes, and our results collectively imply a complex relationship between Cdc14 activity and cell wall integrity signaling in *C. albicans* (see Discussion).

### Cdc14 deficiency impairs *C. albicans* septum assembly and cell separation

We used TEM to look for overt differences in cell wall structure between wild-type and Cdc14-deficient *C. albicans* yeast form cells that could explain the observed cell wall integrity phenotypes. The most striking difference was the structure of the septum in dividing cells. In *CDC14/CDC14* and *CDC14/Δ* cells the primary septum was uniformly visible as a bright, straight band bisecting the bud neck, with thin layers of secondary septum on either side (**Fig 5A-B**), as expected (49). Relatively normal primary septa were also observed in some *cdc14^hm^/Δ* and *cdc14Δ/Δ* cells, however we frequently observed primary septa with abnormal morphologies in these strains that were never observed in wild-type cells (**Fig 5C-D**). The primary septa were often discontinuous, non-linear, forked, or deviated from the bud neck. Moreover, the secondary septa in *cdc14^hm^/Δ* and *cdc14Δ/Δ* were often thickened, and in some cases appeared to lack primary septa altogether. These secondary septa were reminiscent of previously described remedial septa that form in the absence of primary septum assembly (50). From the TEM analysis we conclude that high Cdc14 activity is critical for proper septum assembly during cell division.

**Figure 5.**
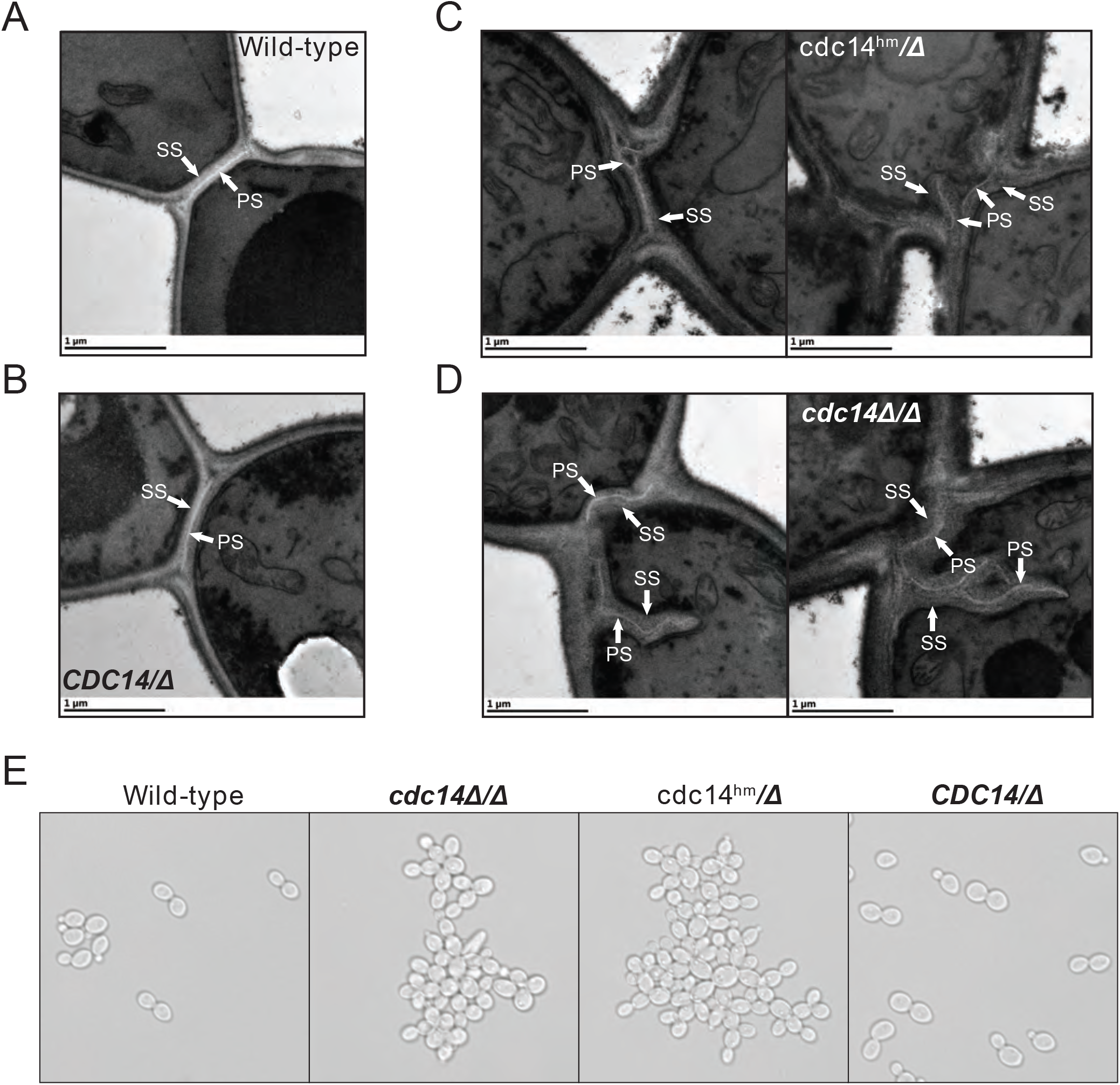
Reduced Cdc14 activity impairs *C. albicans* septum assembly and cell separation. (**A-D**) Transmission electron micrographs of *C. albicans* yeast form cells of the indicated strains undergoing septation in mid-log phase liquid cultures. PS = primary septum, SS = secondary septum. (**E**) Differential interference contrast (DIC) microscopy images of log phase cultures of the indicated strains.

Previous characterization of *cdc14Δ/Δ C. albicans* revealed defects in separation of yeast form cells following cell division, resulting in formation of large cell clumps (29). This phenotype was linked to regulation of the Ace2 transcription factor that controls expression of cell separation genes following cytokinesis and septation. In liquid cultures *cdc14^hm^/Δ* also formed large cell clumps that were indistinguishable from *cdc14Δ/Δ,* whereas *CDC14/Δ* did not (**Fig 5E**). The clumps were dispersed by sonication, consistent with normal completion of cytokinesis, but incomplete dissolution of septa. In asynchronously growing cultures we detected similar reductions in the transcript levels of Ace2 target genes in *cdc14^hm^/Δ* and *cdc14Δ/Δ* by qRT-PCR (**Fig S6**). This suggests that high Cdc14 activity is also needed to fully activate Ace2. Thus, CaCdc14 appears to play critical roles in both the assembly and dissolution of the septum during yeast form cell division. Septum defects could, in principle, contribute to the impaired cell wall integrity of Cdc14-deficient cells.

### Reduced Cdc14 activity impairs *C. albicans* hyphal development and pathogenesis

*C. albicans cdc14Δ/Δ* was also previously reported to be defective in hyphal differentiation and invasive growth on agar plates (29). We therefore tested if *cdc14^hm^/Δ* also exhibits these phenotypes. On Spider medium plates, *cdc14Δ/Δ* and *cdc14^hm^/Δ* were completely unable to develop filamentous colonies and exhibited very limited agar invasion (**Fig 6A**). Microscopic analysis of cells extracted from the Spider agar after washing of the plate surface revealed that even invasive cells exhibited a severe reduction in the number and length of hyphae compared to strains with wild-type *CDC14* (**Fig 6B**). Similar lack of invasion and formation of hyphae by *cdc14Δ/Δ* and *cdc14^hm^/Δ* were also observed on YPD-serum agar plates (**Fig S7A-B**). In liquid cultures, hyphal differentiation differences between the strains were more subtle. Both *cdc14Δ/Δ* and *cdc14^hm^/Δ* exhibited hyper-polarized growth consistent with hyphae, but at two hours after serum induction their elongated cells were shorter on average and displayed more pseudohyphal character than those of the wild-type *CDC14* strains (**Fig S7C**). We conclude that normal hyphal development also requires high *CDC14* activity.

**Figure 6.**
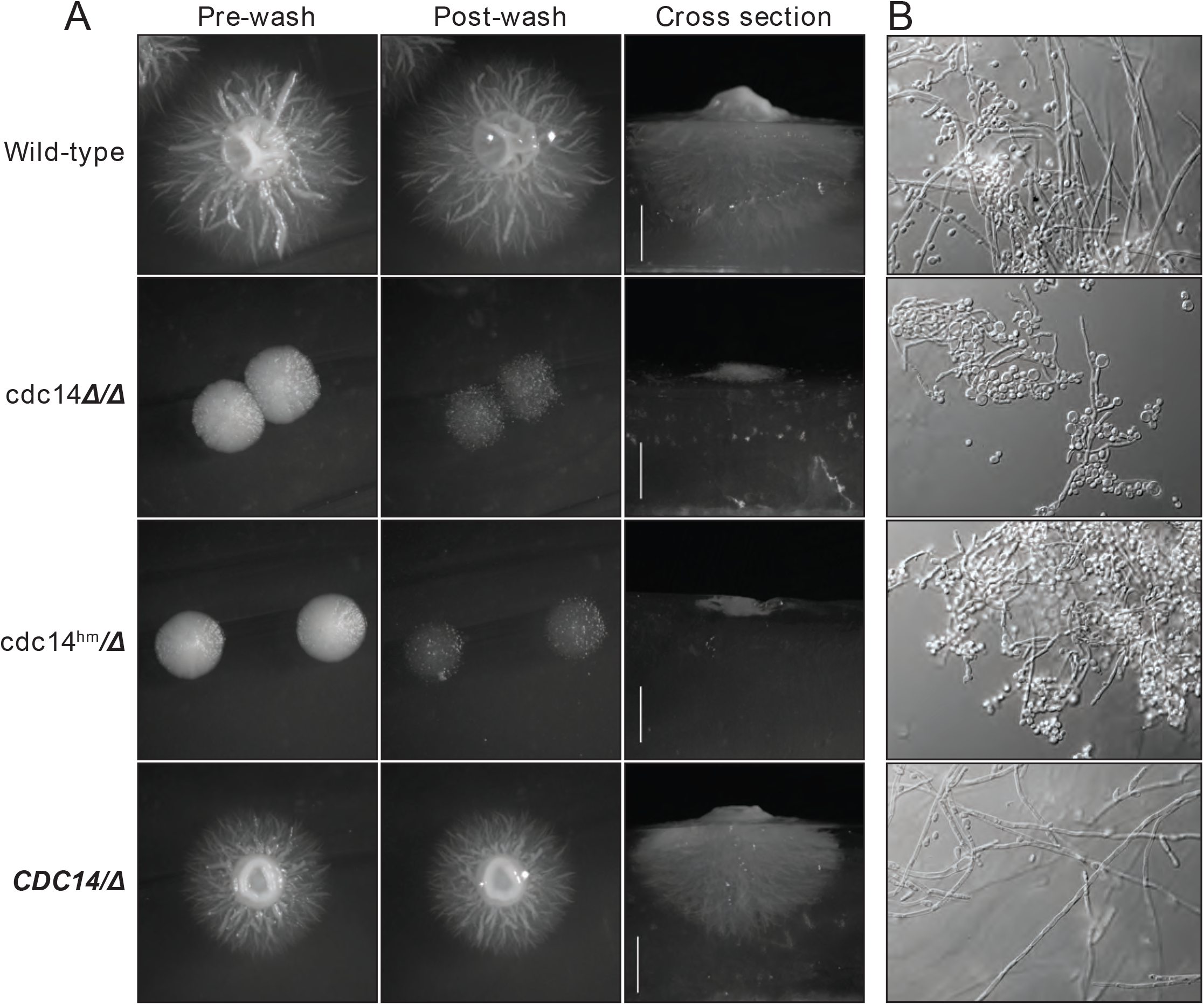
Reduced Cdc14 activity impairs hyphal development on solid medium. (**A**) The indicated *C. albicans* strains were grown on Spider medium agar plates at 37 °C for 5 days. Plates were imaged before and after vigorous washing of unattached cells off the surface with water. After washing, agar plus were removed and cross sections were imaged to illustrate depth of invasive growth. (**B**) Pieces of the agar plugs from (A) were manually ground into small pieces, which were then mounted on a microscopy slide, compressed under a cover slip, and images of embedded cells captured by DIC microscopy.

Hyphal differentiation is thought to be important for *C. albicans* virulence (32), suggesting that reduced Cdc14 activity might compromise pathogenesis. We tested this idea in two established animal models of invasive candidiasis, the larvae of the wax moth *Galleria mellonella* (51–54), and immunosuppressed mice (55). In *G. mellonella, cdc14Δ/Δ* and *cdc14^hm^/Δ* exhibited dramatically reduced virulence compared to *CDC14/CDC14* and *CDC14/Δ* (**Fig 7A**). In mice the virulence defect associated with reduced Cdc14 was even more dramatic. An initial titration of fungal dose revealed *cdc14Δ/Δ* to be almost completely avirulent even at the highest fungal titer (**Fig S8A-B**). At the minimum *CDC14/CDC14* dose required for complete mortality, *cdc14^hm^/Δ* failed to kill any mice and, surprisingly, *CDC14/Δ* resulted in 50% mortality, killing only the female mice, suggesting *CDC14* heterozygosity is sufficient to impact virulence and that there may be a gender bias in sensitivity to Cdc14 deficiency (**Fig 7B**).

**Figure 7.**
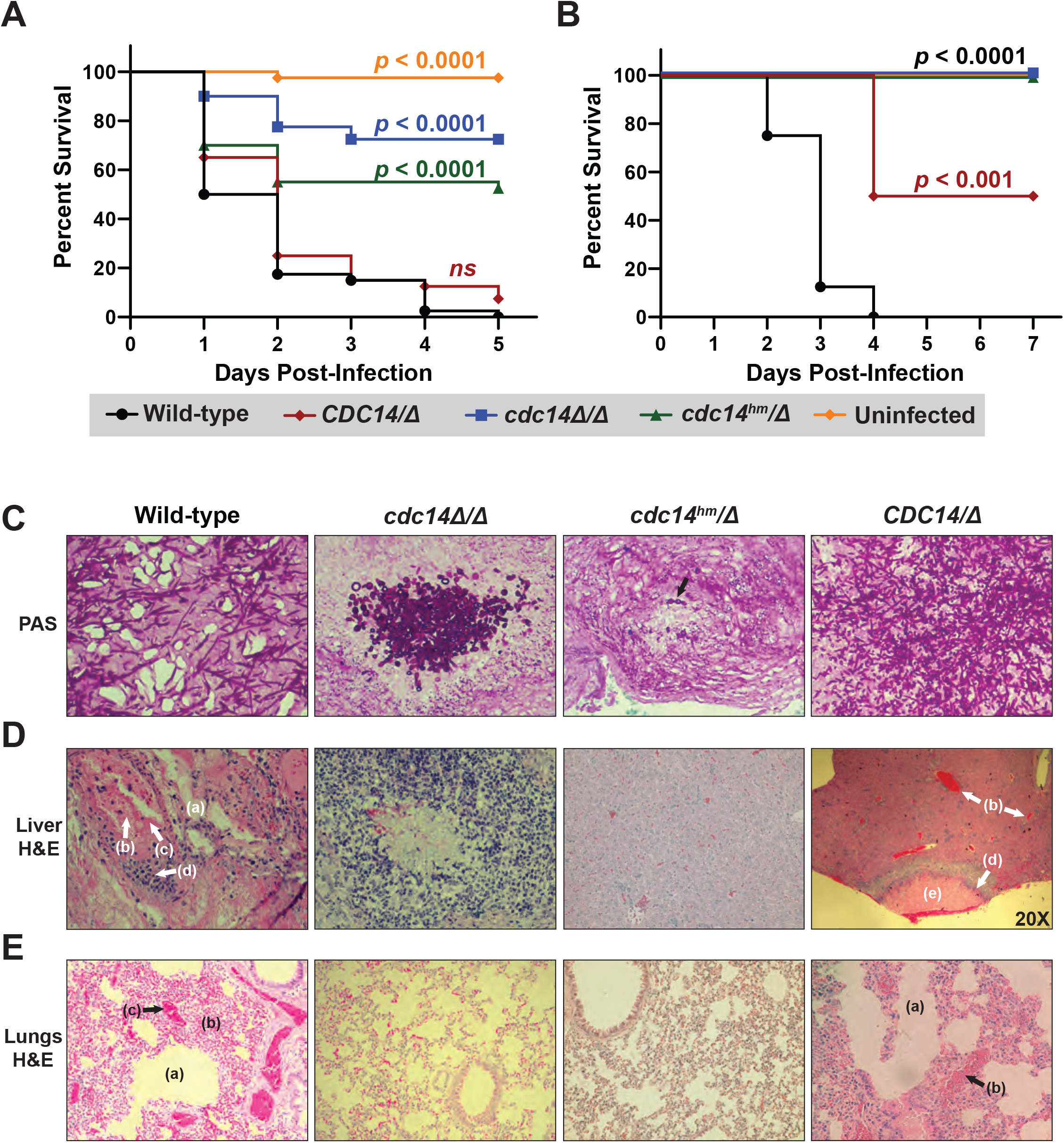
Loss or reduction of Cdc14 function impairs pathogenesis of *C. albicans*. (**A**) *G. mellonella* larvae or (**B**) immunosuppressed BALB/c mice were infected with 2 ×10^5^ or 5 × 10^5^ *C. albicans* cells of the indicated strains, respectively, or mock-uninfected. *p* values were determined using a log-rank (Mantel-Cox) test comparing the survival distribution of hosts infected with wild-type *C. albicans* to each experimental strain. (**C-E**) Histopathological analysis of tissue sections from mouse organs harvested after completion of experiment in (**B**). In (**C**) fungal cells were stained with periodic acid-Schiff (PAS) in representative tissue sections to illustrate morphology. In (**D**) and (**E**) liver and lung tissues, respectively, were stained with H&E. All images were acquired at 200x magnification, except the *CDC14/Δ* sample in (**D**), which was imaged at 20x to illustrate the necrotic lesion with surrounding inflammation. The *cdc14Δ/Δ* liver section in (**D**) illustrates one of the rare lesions (associated with the cluster of fungal cells shown in (**C**)) found in mice infected with *cdc14* mutant strains. In (**D**) and (**E**) the following shock-related lesions (lacking invading fungal cells) in mice infected with *CDC14/CDC14* and *CDC14/Δ* strains are labeled: endothelial leakage characterized by hepatic (**D**) or pulmonary alveolar (**E**) edema (a), multifocal hemorrhage and congestion (b), and intravascular microthrombi (c). Recruited inflammatory cells (d) and necrotic lesions (e) are also labeled. H&E-stained liver and lung sections of an uninfected mouse can be found in **Fig. S8C** for comparison.

Histological analysis of mouse organs following infection revealed extensive colonization by predominantly hyphal cells of *CDC14/CDC14* and *CDC14/Δ* strains, resulting in inflammation and necrosis, as expected (**Fig 7C-E and Fig. S8C**). In contrast, minimal colonization and necrosis were observed after infection by *cdc14^hm^/Δ* and *cdc14Δ/Δ* strains and the rare fungal cells that were detected were almost completely yeast form, consistent with the hyphal differentiation defects observed on agar plates. In mice infected with wild-type *C. albicans,* organs like the liver and lungs showed evidence of shock prior to appearance of fungal cells. Lesions consistent with shock included endothelial leakage characterized by edema fluid, multifocal hemorrhage and congestion, and occasional intravascular microthrombi (**Fig 7D-E**). These were not observed in the organs of mice infected with *cdc14^hm^/Δ* or *cdc14Δ/Δ,* raising the possibility that Cdc14 promotes secretion of soluble factors that contribute to vascular damage. These results show that high Cdc14 activity is critically important for *C. albicans* pathogenesis.

## DISCUSSION

Our results identify Cdc14 phosphatase as a novel virulence factor in *C. albicans* and reveal a new and conserved contribution of Cdc14 to fungal cell wall integrity. Importantly, partial loss of Cdc14 activity through mutation of a conserved catalytic enhancer element in the disordered C-terminal region was sufficient to render *C. albicans* hypersensitive to cell wall stress, and impair septation, hyphal development, and pathogenesis. Since *C. albicans* virulence depends on hyphal differentiation (31,32), the pathogenesis phenotype could be explained by the hyphal defect of Cdc14-deficient strains. However, CaCdc14 function in septation and cell wall integrity may contribute to virulence as well, a possibility that will require future testing. We speculate that the observed cell wall integrity and septation defects reflect a general importance of high level Cdc14 expression across fungi.

### Cdc14 importance for fungal septation

Cdc14 has been implicated in regulation of septation and cell separation previously. In *S. cerevisiae* Cdc14 promotes delivery of the chitin synthase Chs2, which synthesizes the primary septum, to the bud neck after completion of mitosis (56). It also regulates actomyosin ring assembly (57) and the ingression progression complex that coordinates actomyosin ring contraction and membrane ingression with Chs2-dependent septum assembly (58–60). Mutations in the Mitotic Exit Network, which activates ScCdc14, lead to septum structural defects (60). In the phytopathogenic fungi *Fusarium graminearum, Magnaporthe oryzae,* and *Aspergillus flavus, CDC14* deletion reduces the number of septa in conidia and vegetative hyphae and in *M. oryzae* perturbs correct localization of appressoria-delimiting septa (16–18). Despite the evidence that Cdc14 is important for normal septation, our results appear to be the first to show that Cdc14 deficiency causes structural defects in primary septa. Cdc14 is also important for dissolving septa to allow cell separation. ScCdc14 dephosphorylates and activates Ace2 to induce cell separation genes (61,62), and preventing ScCdc14 nuclear export or reducing Cdc14 level via inducible degradation resulted in cell separation failure (22,28). Cdc14 is also needed to activate Ace2 to promote cell separation in *C. albicans* (29). Septation and cell separation defects could be a primary cause of the cell wall stress sensitivity observed in Cdc14-deficient *C. albicans,* although direct evidence of such a causal relationship is still missing.

### A novel role for fungal Cdc14 in cell wall integrity and stress signaling

In addition to causing a cell wall structural defect, our results suggest that Cdc14 deficiency impacts CWI signaling in *C. albicans* in a complex way. Despite the elevated CWI pathway activation in unstressed Cdc14-deficient strains (including heterozygous *CDC14/Δ), cdc14Δ/Δ* and *cdc14^hm^/Δ* exhibited reduced stimulation of phospho-Mkc1 following micafungin treatment. This suggests that Cdc14 deficiency both causes cell wall damage and also plays a role in activating the response to this damage. Consistent with this idea, Cdc14-deficient strains displayed modest up-regulation of a subset of known cell wall stress-responsive genes in the absence of added stress (in line with elevated phospho-Mkc1) but were defective in stimulating expression of these genes in response to micafungin exposure. The extent of this defect correlated with the severity of Cdc14 deficiency; *cdc14Δ/Δ* showed no micafungin-stimulated expression of the four cell wall stress-responsive genes we tested. These results are somewhat difficult to reconcile but are most consistent with Cdc14 influencing CWI signaling at steps upstream and downstream of Mkc1. Consequently, the cell wall stress sensitivity phenotype could be explained by a Cdc14-dependent cell wall structural defect, a deficiency in the CWI signaling response to the cell wall defect, or both. The fact that *CDC14/Δ* cells exhibited increased basal phospho-Mkc1 but normal CWI signaling and transcriptional response after micafungin treatment reinforces the idea that Cdc14 functions in multiple capacities to promote cell wall integrity that are differentially sensitive to its activity level. We surveyed a small subset of cell wall stress-responsive genes, and a complete picture of how Cdc14 impacts the response to cell wall stress will ultimately require global approaches like RNA-seq.

Cdc14 was previously linked to several fungal stress pathways. *S. pombe* Clp1 and ScCdc14 are released from the nucleolus and activated in response to genotoxic stress (63–65). A global phosphatase and kinase interaction network analysis revealed multiple interactions of ScCdc14 with stress signaling MAP kinases (38) and other studies have reported connections between Cdc14 and the osmotic stress response mediated by the Hog1 MAP kinase (66,67). In the phytopathogen *A. flavus,* a *CDC14Δ* strain exhibited growth delays in the presence of chitin-binding compounds and osmotic stress (18). In the entomopathogen *Beauveria bassiana, CDC14Δ* displayed modest sensitivity to multiple stresses, including the chitin-binding Congo red (19). These studies imply that fungal Cdc14 may play diverse roles in facilitating fungal stress responses, although we emphasize that *cdc14^hm^/Δ C. albicans* with partial loss of Cdc14 activity exhibited a very specific sensitivity to cell wall stress. An area for future investigation will be determining if the cell wall stress hypersensitivity of Cdc14-deficient strains is related to the virulence defect. Specific mutations that affect Cdc14 function in cell wall integrity would facilitate this and could be enabled by a thorough understanding of Cdc14 regulation during cell division, in response to cell wall stress, and during hyphal development.

### The significance of Cdc14 activity level and the C-terminal catalytic enhancer motif

Our results reinforce the fact that different functions of Cdc14 require different levels of activity. In *S. cerevisiae* the essential mitotic exit function requires a small fraction of normal Cdc14 activity (22). Why would functions in cytokinesis and cell wall integrity require much higher Cdc14 activity? In *S. cerevisiae* the essential function is fulfilled by nuclear Cdc14 (28) whereas cytokinesis functions require cytoplasmic Cdc14 generated by the Mitotic Exit Network kinases (68). One possibility is that cytoplasmic functions require more enzyme because of the greater volume relative to the nucleus, which effectively dilutes Cdc14 concentration. In *C. albicans* Cdc14 localization is also cell-cycle regulated. It is primarily nuclear prior to mitosis, then appears at the spindle pole bodies during early mitosis, and finally becomes cytoplasmic and localizes to the bud neck in late mitosis (29). A second possibility is that the mitotic exit function requires minimal activity to trigger a positive feedback system (69), whereas the cytokinesis, septation, and cell wall integrity functions depend on bulk substrate dephosphorylation that necessitates higher Cdc14 concentration. A third possibility is that Cdc14 has high affinity for mitotic exit substrates that must be dephosphorylated relatively early after Cdc14 activation and lower affinity for substrates involved in later processes like cytokinesis and septation, resulting in a need for higher Cdc14 activity to achieve their dephosphorylation. There is evidence that substrate affinity does influence the order of Cdc14 substrate dephosphorylation during mitotic exit (70), providing some support for this possibility.

Regardless of the true explanation, it appears clear that Cdc14 activity levels in *C. albicans* and *S. cerevisiae* are physiologically important. Even heterozygous *CDC14/CDC14Δ* cells exhibited some phenotypes, most notably the reduction in virulence in the mouse infection assay. We speculate that the invariant QPRK motif in the C-terminus of fungal Cdc14 enzymes exists to provide a mechanism for regulating Cdc14 activity level around a critical threshold. Although the need for this mechanism is still unclear, results from both *S. cerevisiae* and *S. pombe* support its existence. A C-terminal Cdk phosphorylation site in ScCdc14 was previously shown to inhibit enzyme activity by an unknown mechanism (33). This site happens to be adjacent to the QPRK motif and also appears to be essentially invariant in fungi (although its exact position relative to QPRK varies, see alignment in Fig. 1D). In *S. pombe* Clp1 phosphorylation by Cdk1 at C-terminal sites (including the one adjacent to its QPRK motif) also reduces enzymatic activity and is required for normal kinetics of mitotic progression (34). Thus, it appears likely that phosphoregulation of QPRK motif function is a mechanism for controlling overall Cdc14 activity level in fungal species. It will be interesting to test if phosphorylation adjacent to the QPRK motif of CaCdc14 also regulates enzyme activity and if dephosphorylation is required to promote CaCdc14 function in septation, hyphal development, and the response to cell wall stress.

### CaCdc14 substrates important for septation, CWI, and hyphal development

While a number of substrates of ScCdc14 and SpCdc14 involved in regulation of cytokinesis and/or septation have been identified (71–74), it remains unknown what substrates of CaCdc14 must be dephosphorylated to promote normal septum assembly, cell wall integrity, and hyphal differentiation. Understanding how Cdc14 influences these processes will require identification of the relevant direct substrates. Evidence suggests that Ace2 is a CaCdc14 substrate likely related to the observed cell separation defects in Cdc14-deficient strains (29). A previous proteomic study of CaCdc14 interacting proteins and candidate substrates did not identify proteins with obvious connections to septum assembly but did identify Ace2, the chitinase Cht4, and a couple components of the actomyosin contractile ring (75), although these were not validated as substrates. Thus, our current knowledge of Cdc14 substrates in *C. albicans* is very limited.

### Cdc14 may have potential as an antifungal drug target

The observation that partial reduction in Cdc14 activity severely impairs pathogenesis in animal models of invasive candidiasis suggests that Cdc14 might be a useful antifungal drug target where even modest therapeutic inhibition of activity could successfully combat infections. Moreover, the fact that reduced Cdc14 activity sensitizes *C. albicans* to echinocandin drugs raises the possibility that Cdc14 inhibitors could be valuable for combating clinical echinocandin resistance. We argued previously that the Cdc14 active site should be amenable to specific inhibitor development (4). High resolution structural information on substrate recognition by the fungal Cdc14 active site is available (7), and despite sharing a common catalytic mechanism with PTPs, Cdc14 possesses a strict and unusual substrate specificity that could be mimicked for effective and highly selective inhibitor development. In animals, deletion of CDC14 has little impact on cell division and development (76–78) and the possibility of successful therapeutic intervention by partial inhibition of Cdc14 activity further minimizes the likelihood of deleterious effects on human cells. The Cdc14 active site region is invariant across the fungal kingdom (4) and therefore inhibitors developed to target CaCdc14 would likely be effective at inhibiting Cdc14 orthologs from many other pathogens. For this reason, it will be useful in the future to determine if Cdc14 is similarly required for infection by other human pathogens.

## METHODS

### Strain and plasmid construction

All plasmids and strains used in this study are listed in Supplementary Tables S1 and S2, respectively. All engineered strains were confirmed by PCR and DNA sequencing. *S. cerevisiae* strain YKA1038 expressing *cdc14^hm^* in W303 background was created using the *delitto perfetto* approach exactly as described (37). The plasmid shuffle *CDC14Δ* strain HCY109 and the pRS314 plasmid expressing *CDC14* from its natural promoter were gifts from Dr. Harry Charbonneau, Purdue University. To replace wild-type *CDC14* in HCY109, cells were transformed with pRS314 expressing either *cdc14^hm^* or *CDC14* followed by selection on 5-FOA to eliminate the URA3-based pRS316-*CDC14* plasmid. *C. albicans* strain JC2712 was created by integration of *CaURA3* at the *NEUT5L* locus in CAI4. JC2711 was created by deletion of both copies of *CDC14* using the URA blaster method (79) followed by integration of *CaURA3* at the *NEUT5L* locus. *C. albicans* complementing strains expressing a single-copy of wild-type *CDC14-3xHA* (JC2721) or *cdc14^hm^-3xHA* (HCA1102) were created by transformation of a *SalI* and *Eco* RV fragment of pJC347 or pHLP661, respectively, into JC8 (CAI4 *cdc14Δ/Δ)* and selecting on SD-URA medium. To create pJC347, the 3’ UTR of *CDC14* (+250 to +536 bp after the STOP codon) was PCR-amplified with the addition of *SacI* and *EcoRV* sites to the 5’ and 3’ ends, respectively, and cloned into *Sac* I and *Eco* RV-digested pFA-URA3 (80) to generate pFA-URA3-CDC14utr. Then, a synthetic DNA with *Sal* I and *BamHI* sites added to the 5’ and 3’ ends, respectively, containing 400bp of *CDC14* promoter, the CDC14-3xHA ORF and a 3-UTR with the first 249 bp after the STOP codon of the *CDC14* gene was cloned into the *Sal* I and *BamHI* sites on pFA-URA3-CDC14utr. To create pHLP661, the *cdc14^hm^* mutation (encoding ^414^QPRK➔^414^AARA substitutions) was generated with the QuikChange Lightning site-directed mutagenesis kit (210518, Agilent) using pJC347 as the template and confirmed by DNA sequencing. *yap1Δ* and *hog1Δ* strains were obtained from the *S. cerevisiae* haploid deletion library (Horizon Discovery). The *yen1Δ mus81Δ S. cerevisiae* strain was a gift from Dr. Lorraine Symington, Columbia University. *S. pombe* strains were a gift from Dr. Kathleen Gould, Vanderbilt University. The *cdc14-3* strain was a gift from Dr. Michael Weinreich, Van Andel Research Institute.

*E. coli* expression plasmids for purification of recombinant Cdc14 enzymes were generated using the Gateway cloning system (Invitrogen). *S. cerevisiae CDC14* coding sequence was amplified by PCR from an existing plasmid. *C. albicans CDC14* coding sequence was amplified from a cDNA preparation to avoid its intron. Both were cloned into the pENTR/D-TOPO entry vector and then transferred to pDEST17 destination vector following the provided instructions for expression with an N-terminal 6xHis tag. Mutations to the QPRK motif of both were generated by QuikChange mutagenesis. Full coding sequences for all plasmids were verified by DNA sequencing.

### Cell culture

Liquid *S. cerevisiae* and *C. albicans* cultures were grown in YPD medium (10 g/L yeast extract, 20 g/L peptone, 20 g/L glucose). YPD was supplemented with 40 mg/L adenine (YPAD) for *S. cerevisiae* W303 strains. Cultures were grown at 30°C with shaking at 225 rpm. *S. pombe* strains were grown in YES (5 g/L yeast extract, 30 g/L glucose, 225 mg/L adenine, histidine, leucine, uracil, and lysine) at 30°C with shaking at 225 rpm. *E. coli* were grown in 2xYT (16 g/L tryptone, 10 g/L yeast extract, and 5 g/L NaCl) at 37°C with shaking at 225 rpm. Agar was added to 20% (w/v) for growth on solid medium. For agar plate spotting assays, single colonies were grown to saturation. Strains were serially diluted in 8-fold steps starting from OD_600_ = 1.0 and 5 μL of four consecutive dilutions were spotted on plates and grown at 30°C for 3-5 days. *C. albicans cdc14Δ/Δ* cultures were diluted one less time than other strains and spots therefore correspond to 8-fold higher absorbance; this was necessary to normalize colony density on untreated control plates. For hyphal induction on solid medium, *C. albicans* strains were streaked for single colonies on Spider media plates (10 g/L beef broth, 10 g/L mannitol, 2 g/L K_2_HPO_4_, 20 g/L agar) or YPD plates supplemented with 10% fetal bovine serum (FBS) (Atlanta Biologicals, S11550) and grown at 37°C for 5 days. For liquid hyphal induction, strains were grown overnight to saturation at 30 °C and diluted to an OD_600_ = 0.1 in pre-warmed 37 °C YPD containing 10% FBS. Cultures were then incubated at 37°C with shaking at 225 rpm.

### Immunoblotting

Total protein extracts were prepared as described (81,82) with subtle modifications. Briefly, 8 mL of mid-log phase cells were treated with 10% trichloroacetic acid and pelleted by centrifugation. Cell pellets were washed with 10 mL 70% ethanol and 2x with 1 mL water, then re-suspended in 1 mL 0.2 M NaOH and incubated 10 min on ice. Cells were pelleted, resuspended in 160 x OD_600_ μl loading dye (120 mM Tris-HCl pH 6.8, 4% sodium dodecyl sulfate, 0.02% bromophenol blue, 20% glycerol), heated at 95°C for 10 min, and lysates clarified by centrifugation at 16,000 *x g.* Proteins were separated on 10% tris-glycine SDS-PAGE gels, transferred to 0.45 μm nitrocellulose membranes (Bio-Rad) and probed overnight at 4 °C with mouse anti-HA (1:5,000; Sigma-Aldrich, 12CA5), rabbit anti-PSTAIR (1:5,000; Millipore-Sigma, 06-923), rabbit anti-p44/42 MAPK (1:2,500; Cell Signaling Technology, 9102) or rabbit anti-G6PDH (1:5,000; Sigma-Aldrich, A9521). Secondary anti-mouse and anti-rabbit antibodies conjugated to horseradish peroxidase were from Jackson ImmunoResearch (115-035-003 or 111-035-003) and used at 1:10,000 dilution for 60 min at 4 °C. Immunoblots were developed using Clarity Western ECL Substrate (Bio-Rad, 170-5060) and imaged on a ChemiDoc MP multimode imager (Bio-Rad).

### Protein purification

6xHis-Cdc14 enzymes were purified from *E. coli* as previously described (4).

### Enzyme kinetics

Activities towards varying concentrations of pNPP and synthetic phosphopeptides were assayed as previously described (4).

### qRT-PCR

Cells were grown to mid-log phase (OD_600_ ~ 0.8) in YPD and, where indicated, treated with 50 ng/mL micafungin for 1 hour prior to RNA extraction. RNA was isolated and purified using acid phenol as described (83,84). Reverse transcription was performed using All-in-One 5X RT MasterMix (ABM, G592). Primer Express 3.0 software was used to design primers and qRT-PCR was performed as previously described (83,84) with at least three biological replicates. Data were analyzed using the comparative *C_T_* method (2^-ΔΔ*CT*^). Internal controls were *RDN18* (18S rRNA) for *C. albicans* and *ACT1* for *S. cerevisiae*. All samples were normalized to an untreated wild-type strain.

### Galleria mellonella infection assays

Larvae of *G. mellonella* were purchased from waxworms.net (St. Marys, OH) and infection assays were performed similar to previous reports (52,85). Upon arrival, larvae were held at room temperature in the dark, without food for 24 hours before injection. *C. albicans* strains were grown to saturation in YPD and cells washed three times with PBS, sonicated for 10 seconds, and counted using a Luna-FL automated cell counter (Logos Biosystems). Larvae (n ≥ 40 per strain) of similar size and color were injected with 6 μL of 2×10^5^ cells into their lower left proleg using a Hamilton syringe. An additional group was injected with PBS to serve as a negative control. Injected larvae were incubated in petri dishes at 37°C in the dark for 5 days. Death was measured every 24 hours based on the ability of the larvae to flip over after being placed on their backs. Survival data was plotted using the Kaplan-Meier survival curve and statistical analysis was performed using a log-rank Mantel-Cox test.

### Mouse infection assay

BALB/c mice (n=8 per strain, 4 males and 4 females, 5-week-old) were acclimatized for 7 days upon arrival. On days 8 and 11, mice were rendered neutropenic by intraperitoneal (IP) doses of cyclophosphamide; 150 mg/kg on day 8 and 100 mg/kg on day 11. On day 12, 5 × 10^5^ CFUs of a *C. albicans* strain (measured using a hemocytometer after staining with 0.4% trypan blue to rule out dead cells) in 0.2 mL PBS were administered via IP injection. An additional group received cyclophosphamide but was not infected to serve as a negative control. Mice were monitored for morbidity every 6 hours and were euthanized upon showing severe signs of illness, including weight loss (>20%), dyspnea, unresponsiveness, staggered gait, and inability to eat. On day 19, surviving mice were humanely euthanized through CO2 inhalation. Lungs, liver, spleen, and kidneys were collected from euthanized mice for histopathology. Survival data were plotted using the Kaplan-Meier survival curve and statistical analysis was performed using a log-rank Mantel-Cox test. Selection of 5 × 10^5^ CFUs of each *C. albicans* strain was based on a pilot study performed using different doses (5 × 10^6^ to 1 × 10^4^ CFUs) of both wild-type and *cdc14Δ/Δ* strains (**Fig S8A-B**).

### Histopathology

Lungs, kidneys, and livers from euthanized mice were formalin-fixed, and transferred to 70% ethanol on the next day. Tissues were embedded in paraffin, sectioned, and stained with hematoxylin and eosin (H&E) at the Purdue Histology Research Laboratory for histopathological examination. Tissues were examined by a board-certified veterinary anatomic pathologist, histological features were evaluated, and representative tissue sections were photographed. Fungal pathogens were visualized and evaluated for morphology using periodic acid Schiff (PAS) staining.

### Transmission electron microscopy

Samples were prepared based on a previously published protocol (86) with minor modifications. Day 1: Samples were fixed by adding cell culture to an equal volume of 5% glutaraldehyde in 0.1 M sodium cacodylate buffer. After five min cells were pelleted by centrifugation, resuspended in 2.5% glutaraldehyde in the same buffer, and fixed overnight at 4 °C. Day 2: Cells were rinsed and resuspended in 2% aqueous potassium permanganate for one hour, then rinsed until the solution cleared. Cells were then embedded in agarose, *en bloc* stained in 1% aqueous uranyl acetate for one hour and rinsed. Day 3: Samples were dehydrated with a graded series of ethanol, then transferred into a 2:1 mixture of ethanol and LR White resin and rotated for 2 hours. They were then transferred into a 1:1 mixture of ethanol and LR White resin and rotated overnight. Day 4: Samples were moved into pure LR White resin, rotated for two hours, then placed under house vacuum. The resin was changed three times throughout the day and the samples remained under vacuum overnight. Day 5: Samples were embedded in LR White resin in gelatin capsules and polymerized at 60 °C for 24 hours. Thin sections were cut on a Leica UC6 ultramicrotome and stained with 4% uranyl acetate and lead citrate. Images were acquired on a FEI Tecnai T12 electron microscope equipped with a tungsten source operated at 80 kV.

## ACKNOWLEDGEMENTS

We thank Dr. Kathleen Gould (Vanderbilt University), Dr. Harry Charbonneau (Purdue University), Dr. Michael Weinreich (Van Andel Research Institute), and Dr. Lorraine Symington (Columbia University) for strains and plasmids. We thank Dr. Chris Gilpin and Laurie Muller of the Purdue Life Sciences Microscopy Facility for assistance preparing TEM samples and collecting TEM data. We also thank the Purdue University Animal Resource Facility and Histology Research laboratory for their contributions to this research.

## REFERENCES

1. Culotti J, Hartwell LH. Genetic control of the cell division cycle in yeast. 3. Seven genes controlling nuclear division. Exp Cell Res. 1971 Aug;67(2):389–401.

2. Visintin R, Craig K, Hwang ES, Prinz S, Tyers M, Amon A. The phosphatase Cdc14 triggers mitotic exit by reversal of Cdk-dependent phosphorylation. Mol Cell. 1998 Dec;2(6):709–18.

3. Kerk D, Templeton G, Moorhead GB. Evolutionary radiation pattern of novel protein phosphatases revealed by analysis of protein data from the completely sequenced genomes of humans, green algae, and higher plants. Plant Physiol. 2008 Feb;146(2):351–67.

4. DeMarco AG, Milholland KL, Pendleton AL, Whitney JJ, Zhu P, Wesenberg DT, et al. Conservation of Cdc14 phosphatase specificity in plant fungal pathogens: implications for antifungal development. Sci Rep. 2020 Dec;10(1):12073.

5. Ah-Fong AM, Judelson HS. New role for Cdc14 phosphatase: localization to basal bodies in the oomycete phytophthora and its evolutionary coinheritance with eukaryotic flagella. PLoS One. 2011 Feb 14;6(2):e16725.

6. Gray CH, Good VM, Tonks NK, Barford D. The structure of the cell cycle protein Cdc14 reveals a proline-directed protein phosphatase. EMBO J. 2003 Jul 15;22(14):3524–35.

7. Kobayashi J, Matsuura Y. Structure and dimerization of the catalytic domain of the protein phosphatase Cdc14p, a key regulator of mitotic exit in Saccharomyces cerevisiae. Protein Sci. 2017 Jul 30;26(10):2105–12.

8. Bremmer SC, Hall H, Martinez JS, Eissler CL, Hinrichsen TH, Rossie S, et al. Cdc14 phosphatases preferentially dephosphorylate a subset of cyclin-dependent kinase (Cdk) sites containing phosphoserine. J Biol Chem. 2012 Jan 13;287(3):1662–9.

9. Li L, Ernsting BR, Wishart MJ, Lohse DL, Dixon JE. A family of putative tumor suppressors is structurally and functionally conserved in humans and yeast. J Biol Chem. 1997 Nov 21;272(47):29403–6.

10. Vázquez-Novelle MD, Esteban V, Bueno A, Sacristán MP. Functional Homology among Human and Fission Yeast Cdc14 Phosphatases. J Biol Chem. 2005 Aug 12;280(32):29144–50.

11. Taylor GS, Liu Y, Baskerville C, Charbonneau H. The activity of Cdc14p, an oligomeric dual specificity protein phosphatase from Saccharomyces cerevisiae, is required for cell cycle progression. J Biol Chem. 1997 Sep 19;272(38):24054–63.

12. Wang WQ, Bembenek J, Gee KR, Yu H, Charbonneau H, Zhang ZY. Kinetic and Mechanistic Studies of a Cell Cycle Protein Phosphatase Cdc14. J Biol Chem. 2004;279(29):10.

13. Clement A, Solnica-Krezel L, Gould KL. The Cdc14B phosphatase contributes to ciliogenesis in zebrafish. Development. 2011 Jan;138(2):291–302.

14. Uddin B, Partscht P, Chen NP, Neuner A, Weiß M, Hardt R, et al. The human phosphatase CDC14A modulates primary cilium length by regulating centrosomal actin nucleation. EMBO Rep. 2019 Jan 1;20(1):e46544.

15. Imtiaz A, Belyantseva IA, Beirl AJ, Fenollar-Ferrer C, Bashir R, Bukhari I, et al. CDC14A phosphatase is essential for hearing and male fertility in mouse and human. Hum Mol Genet. 2018 Mar 1;27(5):780–98.

16. Li C, Melesse M, Zhang S, Hao C, Wang C, Zhang H, et al. FgCDC14 regulates cytokinesis, morphogenesis, and pathogenesis in Fusarium graminearum. Mol Microbiol. 2015 Nov;98(4):770–86.

17. Li C, Cao S, Zhang C, Zhang Y, Zhang Q, Xu JR, et al. MoCDC14 is important for septation during conidiation and appressorium formation in Magnaporthe oryzae. Mol Plant Pathol. 2018;19(2):328–40.

18. Yang G, Hu Y, Fasoyin OE, Yue Y, Chen L, Qiu Y, et al. The Aspergillus flavus phosphatase CDC14 regulates development, aflatoxin biosynthesis and pathogenicity. Front Cell Infect Microbiol. 2018 May;8(141).

19. Wang J, Liu J, Hu Y, Ying SH, Feng MG. Cytokinesis-required Cdc14 is a signaling hub of asexual development and multi-stress tolerance in Beauveria bassiana. Sci Rep. 2013 30/online;3:3086.

20. Trautmann S, Wolfe BA, Jorgensen P, Tyers M, Gould KL, McCollum D. Fission yeast Clp1p phosphatase regulates G2/M transition and coordination of cytokinesis with cell cycle progression. Curr Biol. 2001 Jun 26;11(12):931–40.

21. Cueille N, Salimova E, Esteban V, Blanco M, Moreno S, Bueno A, et al. Flp1, a fission yeast orthologue of the s. cerevisiae CDC14 gene, is not required for cyclin degradation or rum1p stabilisation at the end of mitosis. J Cell Sci. 2001 Jul;114(Pt 14):2649–64.

22. Powers BL, Hall MC. Re-examining the role of Cdc14 phosphatase in reversal of Cdk phosphorylation during mitotic exit. J Cell Sci. 2017 Jan 1;jcs.201012.

23. Jungbluth M, Renicke C, Taxis C. Targeted protein depletion in Saccharomyces cerevisiae by activation of a bidirectional degron. BMC Syst Biol. 2010;4:176.

24. Taxis C, Stier G, Spadaccini R, Knop M. Efficient protein depletion by genetically controlled deprotection of a dormant N-degron. Mol Syst Biol. 2009;5:267.

25. Wang M, Herrmann CJ, Simonovic M, Szklarczyk D, Mering C. Version 4.0 of PaxDb: Protein abundance data, integrated across model organisms, tissues, and cell-lines. Proteomics. 2015 Sep;15(18):3163–8.

26. Manzano-López J, Monje-Casas F. The Multiple Roles of the Cdc14 Phosphatase in Cell Cycle Control. Int J Mol Sci. 2020;21(3).

27. Clifford DM, Wolfe BA, Roberts-Galbraith RH, McDonald WH, Yates JR, Gould KL. The Clp1/Cdc14 phosphatase contributes to the robustness of cytokinesis by association with anillin-related Mid1. J Cell Biol. 2008 Apr 7;181(1):79–88.

28. Bembenek J, Kang J, Kurischko C, Li B, Raab JR, Belanger KD, et al. Crm1-mediated nuclear export of Cdc14 is required for the completion of cytokinesis in budding yeast. Cell Cycle. 2005 Jul;4(7):961–71.

29. Clemente-Blanco A, Gonzalez-Novo A, Machin F, Caballero-Lima D, Aragon L, Sanchez M, et al. The Cdc14p phosphatase affects late cell-cycle events and morphogenesis in Candida albicans. J Cell Sci. 2006 Mar 15;119(Pt 6):1130–43.

30. Manzano-López J, Monje-Casas F. The Multiple Roles of the Cdc14 Phosphatase in Cell Cycle Control. Int J Mol Sci. 2020;21(3).

31. Brand A. Hyphal growth in human fungal pathogens and its role in virulence. Naglik JR, editor. Int J Microbiol. 2011 Nov 9;2012:517529.

32. Lo HJ, Köhler JR, DiDomenico B, Loebenberg D, Cacciapuoti A, Fink GR. Nonfilamentous C. albicans mutants are avirulent. Cell. 1997 Sep 5;90(5):939–49.

33. Li Y, Cross FR, Chait BT. Method for identifying phosphorylated substrates of specific cyclin/cyclin-dependent kinase complexes. Proc Natl Acad Sci U A. 2014 Aug 5;111(31):11323–8.

34. Wolfe BA, McDonald WH, Yates JR, Gould KL. Phospho-regulation of the Cdc14/Clp1 phosphatase delays late mitotic events in S. pombe. Dev Cell. 2006 Sep 1;11(3):423–30.

35. Taylor GS, Liu Y, Baskerville C, Charbonneau H. The Activity of Cdc14p, an Oligomeric Dual Specificity Protein Phosphatase from Saccharomyces cerevisiae, Is Required for Cell Cycle Progression. J Biol Chem. 1997 Sep;272(38):24054–63.

36. Eissler CL, Mazon G, Powers BL, Savinov SN, Symington LS, Hall MC. The Cdk/Cdc14 module controls activation of the Yen1 holliday junction resolvase to promote genome stability. Mol Cell. 2014 Apr 10;54(1):80–93.

37. Storici F, Resnick MA. The delitto perfetto approach to in vivo site-directed mutagenesis and chromosome rearrangements with synthetic oligonucleotides in yeast. In: Methods in Enzymology [Internet]. Academic Press; 2006. p. 329–45. Available from: https://www.sciencedirect.com/science/article/pii/S0076687905090191

38. Breitkreutz A, Choi H, Sharom JR, Boucher L, Neduva V, Larsen B, et al. A global protein kinase and phosphatase interaction network in yeast. Science. 2010 May 21;328(5981):1043–6.

39. Sucher AJ, Chahine EB, Balcer HE. Echinocandins: the newest class of antifungals. Ann Pharmacother. 2009 Oct 1;43(10):1647–57.

40. Fuchs Beth Burgwyn, Mylonakis Eleftherios. Our paths might cross: the role of the fungal cell wall integrity pathway in stress response and cross talk with other stress response pathways. Eukaryot Cell. 2009 Nov 1;8(11):1616–25.

41. Torres L, Martín H, García-Saez MI, Arroyo J, Molina M, Sánchez M, et al. A protein kinase gene complements the lytic phenotype of Saccharomyces cerevisiae lyt2 mutants. Mol Microbiol. 1991 Nov;5(11):2845–54.

42. Levin DE. Regulation of cell wall biogenesis in Saccharomyces cerevisiae: the cell wall integrity signaling pathway. Genetics. 2011;189(4):1145–75.

43. Zhao X, Muller EG, Rothstein R. A suppressor of two essential checkpoint genes identifies a novel protein that negatively affects dNTP pools. Mol Cell. 1998;2:329–40.

44. Mazur P, Morin N, Baginsky W, el-Sherbeini M, Clemas J A, Nielsen J B, et al. Differential expression and function of two homologous subunits of yeast 1,3-beta-D-glucan synthase. Mol Cell Biol. 1995 Oct 1;15(10):5671–81.

45. Jung US, Levin DE. Genome-wide analysis of gene expression regulated by the yeast cell wall integrity signalling pathway. Mol Microbiol. 1999 Dec 1;34(5):1049–57.

46. Pérez P, Cortés JCG, Cansado J, Ribas JC. Fission yeast cell wall biosynthesis and cell integrity signalling. Cell Surf. 2018 Dec;4:1–9.

47. Roemer T, Jiang B, Davison J, Ketela T, Veillette K, Breton A, et al. Large-scale essential gene identification in Candida albicans and applications to antifungal drug discovery. Mol Microbiol. 2003;50(1):167–81.

48. Ibe C, Munro CA. Fungal cell wall proteins and signaling pathways form a cytoprotective network to combat stresses. J Fungi. 2021;7(9).

49. Meitinger F, Pereira G. Visualization of cytokinesis events in budding yeast by transmission electron microscopy. In: Sanchez-Diaz A, Perez P, editors. Yeast Cytokinesis: Methods and Protocols [Internet]. New York, NY: Springer New York; 2016. p. 87–95. Available from: https://doi.org/10.1007/978-1-4939-3145-3_7

50. Walker LA, Lenardon MD, Preechasuth K, Munro CA, Gow NAR. Cell wall stress induces alternative fungal cytokinesis and septation strategies. J Cell Sci. 2013 Jun 15;126(12):2668–77.

51. Fuchs BB, O’Brien E, El Khoury JB, Mylonakis E. Methods for using Galleria mellonella as a model host to study fungal pathogenesis. Virulence. 2010 Nov 1;1(6):475–82.

52. Cotter G, Doyle S, Kavanagh K. Development of an insect model for the in vivo pathogenicity testing of yeasts. Pathog Dis. 2000;27(2):163–9.

53. Segal E, Frenkel M. Experimental in vivo models of candidiasis. J Fungi. 2018;4(1).

54. MacCallum DM. Hosting infection: experimental models to assay Candida virulence. Tavanti A, editor. Int J Microbiol. 2011 Dec 22;2012:363764.

55. Hohl TM. Overview of vertebrate animal models of fungal infection. J Immunol Methods. 2014 Aug 1;410:100–12.

56. Chin CF, Bennett AM, Ma WK, Hall MC, Yeong FM. Dependence of Chs2 ER export on dephosphorylation by cytoplasmic Cdc14 ensures that septum formation follows mitosis. Lew D, editor. Mol Biol Cell. 2012 Jan;23(1):45–58.

57. Miller DP, Hall H, Chaparian R, Mara M, Mueller A, Hall MC, et al. Dephosphorylation of Iqg1 by Cdc14 regulates cytokinesis in budding yeast. Mol Biol Cell. 2015 Aug 15;26(16):2913–26.

58. Foltman M, Molist I, Arcones I, Sacristan C, Filali-Mouncef Y, Roncero C, et al. Ingression progression complexes control extracellular matrix remodelling during cytokinesis in budding yeast. PLoS Genet. 2016;12(2):e1005864.

59. Palani S, Meitinger F, Boehm ME, Lehmann WD, Pereira G. Cdc14-dependent dephosphorylation of Inn1 contributes to Inn1-Cyk3 complex formation. J Cell Sci. 2012 Mar 27;

60. Meitinger F, Petrova B, Lombardi IM, Bertazzi DT, Hub B, Zentgraf H, et al. Targeted localization of Inn1, Cyk3 and Chs2 by the mitotic-exit network regulates cytokinesis in budding yeast. J Cell Sci. 2010 Jun 1;123(Pt 11):1851–61.

61. Brace J, Hsu J, Weiss EL. Mitotic exit control of the Saccharomyces cerevisiae Ndr/LATS kinase Cbk1 regulates daughter cell separation after cytokinesis. Mol Cell Biol. 2011 Feb;31(4):721–35.

62. Powers BL, Hall H, Charbonneau H, Hall MC. A Substrate Trapping Method for Identification of Direct Cdc14 Phosphatase Targets. Methods Mol Biol. 2017;1505:119–32.

63. Broadus MR, Gould KL. Multiple protein kinases influence the redistribution of fission yeast Clp1/Cdc14 phosphatase upon genotoxic stress. Mol Biol Cell. 2012 Oct;23(20):4118–28.

64. Villoria MT, Ramos F, Duenas E, Faull P, Cutillas PR, Clemente-Blanco A. Stabilization of the metaphase spindle by Cdc14 is required for recombinational DNA repair. EMBO J. 2017 Jan 4;36(1):79–101.

65. Diaz-Cuervo H, Bueno A. Cds1 controls the release of Cdc14-like phosphatase Flp1 from the nucleolus to drive full activation of the checkpoint response to replication stress in fission yeast. Mol Biol Cell. 2008 Jun;19(6):2488–99.

66. Chasman D, Ho YH, Berry DB, Nemec CM, MacGilvray ME, Hose J, et al. Pathway connectivity and signaling coordination in the yeast stress-activated signaling network. Mol Syst Biol. 2014 Nov 19;10(11):759.

67. Tognetti S, Jimenez J, Vigano M, Duch A, Queralt E, de Nadal E, et al. Hog1 activation delays mitotic exit via phosphorylation of Net1. Proc Natl Acad Sci U A. 2020 Apr 21;117(16):8924–33.

68. Sanchez-Diaz A, Nkosi PJ, Murray S, Labib K. The Mitotic Exit Network and Cdc14 phosphatase initiate cytokinesis by counteracting CDK phosphorylations and blocking polarised growth. EMBO J. 2012 Aug 29;31(17):3620–34.

69. López-Avilés S, Kapuy O, Novák B, Uhlmann F. Irreversibility of mitotic exit is the consequence of systems-level feedback. Nature. 2009 May 1;459(7246):592–5.

70. Kataria M, Mouilleron S, Seo MH, Corbi-Verge C, Kim PM, Uhlmann F. A PxL motif promotes timely cell cycle substrate dephosphorylation by the Cdc14 phosphatase. Nat Struct Mol Biol. 2018 Dec;25(12):1093–102.

71. Kuilman T, Maiolica A, Godfrey M, Scheidel N, Aebersold R, Uhlmann F. Identification of Cdk targets that control cytokinesis. EMBO J. 2015 Jan 2;34(1):81–96.

72. Chen JS, Broadus MR, McLean JR, Feoktistova A, Ren L, Gould KL. Comprehensive proteomics analysis reveals new substrates and regulators of the fission yeast clp1/cdc14 phosphatase. Mol Cell Proteomics. 2013 May;12(5):1074–86.

73. Meitinger F, Palani S, Pereira G. The power of MEN in cytokinesis. Cell Cycle. 2012 Jan 15;11(2):219–28.

74. Mocciaro A, Schiebel E. Cdc14: a highly conserved family of phosphatases with non-conserved functions? J Cell Sci. 2010 Sep 1;123(17):2867–76.

75. Kaneva IN, Sudbery IM, Dickman MJ, Sudbery PE. Proteins that physically interact with the phosphatase Cdc14 in Candida albicans have diverse roles in the cell cycle. Sci Rep. 2019 Apr 18;9(1):6258.

76. Mocciaro A, Berdougo E, Zeng K, Black E, Vagnarelli P, Earnshaw W, et al. Vertebrate cells genetically deficient for Cdc14A or Cdc14B retain DNA damage checkpoint proficiency but are impaired in DNA repair. J Cell Biol. 2010 May 17;189(4):631–9.

77. Berdougo E, Nachury MV, Jackson PK, Jallepalli PV. The nucleolar phosphatase *Cdc14B* is dispensable for chromosome segregation and mitotic exit in human cells. Cell Cycle. 2008 May;7(9):1184–90.

78. Saito RM, Perreault A, Peach B, Satterlee JS, van den Heuvel S. The CDC-14 phosphatase controls developmental cell-cycle arrest in C. elegans. Nat Cell Biol. 2004 Aug 1;6(8):777–83.

79. Fonzi WA, Irwin MY. Isogenic strain construction and gene mapping in Candida albicans. Genetics. 1993 Jul 1;134(3):717–28.

80. Gola S, Martin R, Walther A, Dünkler A, Wendland J. New modules for PCR-based gene targeting in Candida albicans: rapid and efficient gene targeting using 100 bp of flanking homology region. Yeast. 2003 Dec 1;20(16):1339–47.

81. Kushnirov VV. Rapid and reliable protein extraction from yeast. Yeast. 2000 Jun 30;16(9):857–60.

82. von der Haar T. Optimized protein extraction for quantitative proteomics of yeasts. PLOS ONE. 2007 Oct 24;2(10):e1078.

83. Baker KM, Hoda S, Saha D, Gregor JB, Georgescu L, Serratore ND, et al. The Set1 Histone H3K4 Methyltransferase Contributes to Azole Susceptibility in a Species-Specific Manner by Differentially Altering the Expression of Drug Efflux Pumps and the Ergosterol Gene Pathway. Antimicrob Agents Chemother. 2022;66(5):20.

84. Serratore ND, Baker KM, Macadlo LA, Gress AR, Powers BL, Atallah N, et al. A novel sterol-signaling pathway governs azole antifungal drug resistance and hypoxic gene repression in Saccharomyces cerevisiae. Genetics. 2018 Mar;208(3):1037–55.

85. Ames L, Duxbury S, Pawlowska B, Ho H lui, Haynes K, Bates S. Galleria mellonella as a host model to study Candida glabrata virulence and antifungal efficacy. Virulence. 2017 Nov 17;8(8):1909–17.

86. Wright R. Transmission electron microscopy of yeast. Microsc Res Tech. 2000 Dec 15;51(6):496–510.

87. Ho CK, Mazon G, Lam AF, Symington LS. Mus81 and Yen1 promote reciprocal exchange during mitotic recombination to maintain genome integrity in budding yeast. Mol Cell. 2010 Dec 22;40(6):988–1000.

