## Supplementary Material for "Reduced Cdc14 phosphatase activity impairs septation, hyphal differentiation and pathogenesis and causes echinocandin hypersensitivity in *Candida albicans*"

### SUPPLEMENTARY TABLES

#### Table S1. Plasmids

| Species | Name | Backbone | Marker | Expressed Protein | Source |
|---|---|---|---|---|---|
| ***E. coli*** | | | | | |
| | pHLP603 | pDEST17 | Ampicillin | 6xHis-ScCdc14(1-551) | This study |
| | pHLP695 | pDEST17 | Ampicillin | 6xHis-ScCdc14(1-449) | This study |
| | pHLP690 | pDEST17 | Ampicillin | 6xHis-ScCdc14(1-374) | This study |
| | pHLP621 | pDEST17 | Ampicillin | 6xHis-ScCdc14(1-449$^{QARA}$) | This study |
| | pHLP622 | pDEST17 | Ampicillin | 6xHis-ScCdc14(1-449$^{APRK}$) | This study |
| | pHLP683 | pDEST17 | Ampicillin | 6xHis-CaCdc14(1-427) | This study |
| | pHLP687 | pDEST17 | Ampicillin | 6xHis-CaCdc14(1-427$^{AARA}$) | This study |
| | | **Origin** | | | |
| ***S. cerevisiae*** | | | | | |
| | pRS314-*CDC14* | CEN/ARS | *TRP1* | ScCdc14 | (1) |
| | pRS314-*cdc14*$^{hm}$ | CEN/ARS | *TRP1* | ScCdc14$^{QARA}$ | This study |
| ***C. albicans*** | | | | | |
| | pJC347 | Integrating | *URA3* | CaCdc14-3xHA | This study |
| | pHLP661 | Integrating | *URA3* | CaCdc14$^{AARA}$-3xHA | This study |

#### Table S2. Yeast strains

| | Strain Name | Relevant Genotype | Source |
|---|---|---|---|
| ***S. cerevisiae*** | | | |
| | W303 | *MATα ade2-1 can1-100 his3-11,15 leu2-3,112 trp1-1 ura3-1* | |
| | YKA1038 | W303 *MATα cdc14*$^{hm}$ | This study |
| | HCY109 | W303 *MATa cdc14::HIS3* pRS316-*CDC14* | (1) |
| | M609 | W303 *MATa cdc14-3* | (2) |
| | *hog1Δ* | BY4741 *MATa hog1::*KanMX4 | Horizon Discovery |
| | *yap1Δ* | BY4741 *MATa yap1::*KanMX4 | Horizon Discovery |
| | LSY2244-90B | *MATα mus81::*KanMX6 *yen1::HIS3 his3::*HphMX4 | (3) |
| ***S. pombe*** | | | |
| | KLG248 | ura4-D18 ade6-216 leu1-32 h- | (4) |
| | KLG3381 | Clp1::ura4 ura4-D18 ade6-216 leu1-32 h- | (4) |
| ***C. albicans*** | | | |
| | CAI4 | *ura3::imm434/ura3::imm434* | (5) |
| | JC8 | CAI4 *cdc14::hisG/cdc14::hisG* | (6) |
| | JC2712 | CAI4 *NEUT5L::URA3* | This study |
| | JC2711 | CAI4 *NEUT5L::URA3 cdc14::hisG/cdc14::hisG* | This study |
| | JC2721 | CAI4 *CDC14-3HA:URA3/cdc14::hisG* | This study |
| | HCAL102 | CAI4 *cdc14*$^{hm}$*-3HA:URA3/cdc14::hisG* | This study |
| | *MKC1-tet* | CaSS1 *mkc1::HIS3/SAT1-Tet*$_{p}$*-MKC1* | (7) |
| | *CEK1-tet* | CaSS1 *mkc1::HIS3/SAT1-Tet*$_{p}$*-CEK1* | (7) |

**Table S3. Primers for qRT-PCR**

| Primer Name | Sequence (5'-3') |
|---|---|
| ***S. cerevisiae*** | |
| ScACT1-001F | TGGATTCCGGTGATGGTGTT |
| ScACT1-002R | TCAAAATGGCGTGAGGTAGAGA |
| ScFKS2-001F | GATGGGACTGGTGACGGTAATTA |
| ScFKS2-002R | TCATCGTACGCACTTTGATCCT |
| ***C. albicans*** | |
| CaRDN18-001F | GCTGATGACTTGCGCTTACTA |
| CaRDN18-002R | GAGAGGTCTGGGAAATCTTGTG |
| CaECM331-001F | GTTGGTCTCATTACCTTCCT |
| CaECM331-002R | CCAACCCGTTCAAATTAGTC |
| CaPGA13-001F | CCACCACTGCTGAACAACCA |
| CaPGA13-002R | GCAGGAGTAGTAACCACAGATTCAATA |
| CaPGA31-001F | TGGTTCCTCCACTGTCAGACAA |
| CaPGA31-002R | ACCACTACCACCCAATTGAAGAA |
| CaCRH11-001F | TGGTCCGTTGACGGTAGTGTT |
| CaCRH11-002R | TTGTGGGAAACCTTGTGCATT |
| CaENG1-001F | GGGCAACTGGTTTGTTCGTC |
| CaENG1-002R | ACCACCTCTTGCTTCCATCG |
| CaDSE1-001F | AACGAAGATCAACGTCAAAGTGATAA |
| CaDSE1-002R | AACTTCATTTTCAACATCAGTCACAAC |

### SUPPLEMENTARY FIGURE LEGENDS

**Figure S1. Purification of ScCdc14 enzymes**. (**A**) 2 μg of ScCdc14 enzyme variants purified from *E. coli* by nickel affinity chromatography were evaluated by SDS-PAGE with Coomassie blue staining. M = molecular weight markers. Lane labels indicate expressed amino acid regions and any amino acid substitutions made in the QPRK motif. (**B**) Relative rates of ScCdc14(1-449$^{QARA}$)-catalyzed dephosphorylation of the phosphopeptide sequences shown in the table at left were measured under identical conditions at 100 μM each substrate. Peptide 1 is the same near-optimal substrate sequence used in Fig 1C and 1F and the other peptides contain substitutions at positions known to influence substrate recognition by ScCdc14. Activity sharply decreased with changes to phosphoamino acid (pS, pT, pY) and the +1 and +3 positions relative to pS, similar to wild-type Cdc14 enzymes (8,9).

**Figure S2. Sequence alignment of fungal and animal Cdc14 orthologs.** Alignment of selected fungal (light blue) and animal (red) *CDC14* homolog sequences was generated using Clustal Omega with default settings. The location of the fungal-specific QPRK motif is indicated by yellow highlighting and conserved residues in the active site region responsible for catalysis and substrate recognition are labeled.

**Figure S3. Characterization of *cdc14$^{hm}$* allele in *S. cerevisiae*.** (**A**) Microplate growth assay in liquid YPAD performed at 30°C. (**B-E**) *S. cerevisiae* strains were serially diluted and spotted on YPAD agar plates containing indicated concentrations of (**B**) hydroxyurea (HU), (**C**) sodium chloride (NaCl), (**D**) hydrogen peroxide ($H_2O_2$), and (**E**) Congo red (CR). "# 1-3" indicates independent isolates of *cdc14$^{hm}$/Δ* strain. *yen1Δ mus81Δ*, *yap1Δ*, and *hog1Δ* strains are positive controls for sensitivity to HU, NaCl, and $H_2O_2$, respectively. (**F**) The same spotting assays on YPAD plates grown at the indicated temperatures. (**G**) Spotting assays were performed with a *cdc14Δ* plasmid shuffle strain kept alive by CEN plasmids expressing either *cdc14$^{hm}$* or wild-type *CDC14* alleles expressed from the natural *CDC14* promoter. (**H**) Spotting assays with *cdc14$^{hm}$* with and without a CEN plasmid expressing wild-type *CDC14* from its natural promoter, compared to wild-type control. All plates were incubated at 30°C (unless otherwise specified) for 72 hours. Wild-type refers to W303 parent strain. Images are representative of three biological replicates. (**I**) Gene expression (qRT-PCR) analysis of *FKS2* in Wild-type and $cdc14^{hm}$. *FKS2* transcript levels were normalized to *ACT1* and set relative to wild-type. Error bars represent standard deviation of three biological replicates each with three technical repeats. T-test *p* value = 0.064.

**Figure S4. Characterization of $cdc14^{hm}$ mutation in *C. albicans*.** (**A**) 2 µg of truncated (residues 1-427) CaCdc14 enzymes (wild-type and QPRK→AARA mutant) purified from *E. coli* by nickel affinity chromatography were analyzed by SDS-PAGE with Coomassie blue staining. M = molecular weight marker. (**B**) Comparison of reaction velocities of CaCdc14(1-427) and CaCdc14($1\text{-}427^{AARA}$) towards a collection of peptides with substitutions at positions important for optimal Cdc14 substrates. Activity is sharply reduced by changes to phosphoamino acid (pS, pT, pY) or the +1 or +3 positions relative to pS, similar to wild-type Cdc14 enzymes from diverse species (9). (**C**) Anti-HA immunoblotting analysis of *C. albicans cdc14Δ/Δ* complemented with wild-type *CDC14-3xHA* and $cdc14^{hm}$*-3xHA* integration cassettes in the absence and presence of 50 ng/ml micafungin (MF). Anti-PSTAIR is a loading control. "# 1-3" refers to individual isolates of $cdc14^{hm}/\Delta$ strain. (**D**) Growth of the indicated *C. albicans* strains in YPD liquid cultures was monitored by absorbance at 600 nm. Samples were sonicated and diluted to $A_{600}$ <1.5 prior to measurement and the undiluted absorbance equivalents plotted. Average values with standard deviation error bars from 3 independent cultures per strain are plotted. Fit curves were generated in GraphPad Prism using the Gompertz growth function. (**E**) *C. albicans* strains were serially diluted and spotted on YPD plates containing the indicated amounts of sodium chloride (NaCl), methyl methanesulfonate (MMS), or hydrogen peroxide ($H_2O_2$). The concentration of stress agents was the maximum tolerated by the wild-type strain without significant reduction in viability or growth rate. All plates were incubated at 30°C for 72 hours. Images are representative of three independent trials.

**Figure S5. Validation of anti-p44/p42 MAP kinase antibody for monitoring Mkc1 activation.** Tet-repressible *CEK1* and *MKC1* strains were grown to log phase, treated with micafungin (to stimulate Mkc1 activation) or micafungin + doxycycline (to shut off expression) and then immunoblotted with antip44/p42 MAP kinase antibody (pMAPK). In the *MKC1-tet* strain, the signal at ~70 kDa was stimulated by micafungin and then repressed by doxycycline, indicating that this band corresponds to phospho-Mkc1.

**Figure S6.** Relative mRNA levels of selected cell wall stress-responsive genes in mid-log phase cultures of *C. albicans* strains were measured by qRT-PCR. Transcript levels were normalized to *RDN18* and set relative to the wild-type sample. Data are averages of 6 biological trials and error bars are standard deviations. Unpaired T-tests assuming equal variance were used to compare the average expression to that of the wild-type strain (* = $p < 0.05$).

**Figure S7. Hyphal development is impaired in *cdc14*Δ/Δ and *cdc14*$^{hm}$/*cdc14*Δ.** (**A-B**) *C. albicans* strains were streaked out on YPD + 10% fetal bovine serum (FBS) and allowed to grow at 37°C for 5 days. Colonies were imaged before and after washing of the plate surface with water. Embedded cells were excised in small agar plugs and imaged by DIC microscopy using a 40x objective. (**C**) *C. albicans* cells grow in YPD + 10% FBS liquid medium at 37 °C for 2 hours were similarly imaged by DIC microscopy. The length of hyphal projections, defined as cell bodies with a length at least twice the width, were measured using ImageJ (minimum 70 per strain). Statistical significance was determined with unpaired T-tests assuming equal variance comparing each strain to wild-type (* = $p < 0.05$; ** = $p < 0.01$).

**Figure S8. Establishment of immune-compromised mouse model of invasive candidiasis.** Immunosuppressed BALB/c mice were infected with indicated amounts of either wild-type **(A)** or *cdc14Δ/Δ* **(B)** *C. albicans* cells or mock-infected to determine the minimum wild-type dose required for complete mortality and assess initial *CDC14* requirement for virulence. *p* values were determined using a log-rank (Mantel-Cox) test comparing the survival distribution of mice infected with wild-type vs *cdc14Δ/Δ* strains at each fungal cell dose. ns = not significant ($p > 0.05$). **C)** Representative images of liver and lung tissue sections from an uninfected mouse from experiment in Fig. 7B.

### SUPPLEMENTARY REFERENCES


1. Taylor GS, Liu Y, Baskerville C, Charbonneau H. The activity of Cdc14p, an oligomeric dual specificity protein phosphatase from Saccharomyces cerevisiae, is required for cell cycle progression. J Biol Chem. 1997 Sep 19;272(38):24054–63.

2. Miller CT, Gabrielse C, Chen YC, Weinreich M. Cdc7p-Dbf4p Regulates Mitotic Exit by Inhibiting Polo Kinase. PLOS Genet. 2009 May 29;5(5):e1000498.

3. Ho CK, Mazon G, Lam AF, Symington LS. Mus81 and Yen1 promote reciprocal exchange during mitotic recombination to maintain genome integrity in budding yeast. Mol Cell. 2010 Dec 22;40(6):988–1000.

4. Broadus MR, Gould KL. Multiple protein kinases influence the redistribution of fission yeast Clp1/Cdc14 phosphatase upon genotoxic stress. Mol Biol Cell. 2012 Oct;23(20):4118–28.

5. Fonzi WA, Irwin MY. Isogenic strain construction and gene mapping in Candida albicans. Genetics. 1993 Jul 1;134(3):717–28.

6. Clemente-Blanco A, Gonzalez-Novo A, Machin F, Caballero-Lima D, Aragon L, Sanchez M, et al. The Cdc14p phosphatase affects late cell-cycle events and morphogenesis in Candida albicans. J Cell Sci. 2006 Mar 15;119(Pt 6):1130–43.

7. Roemer T, Jiang B, Davison J, Ketela T, Veillette K, Breton A, et al. Large-scale essential gene identification in Candida albicans and applications to antifungal drug discovery. Mol Microbiol. 2003;50(1):167–81.

8. Bremmer SC, Hall H, Martinez JS, Eissler CL, Hinrichsen TH, Rossie S, et al. Cdc14 phosphatases preferentially dephosphorylate a subset of cyclin-dependent kinase (Cdk) sites containing phosphoserine. J Biol Chem. 2012 Jan 13;287(3):1662–9.

9. DeMarco AG, Milholland KL, Pendleton AL, Whitney JJ, Zhu P, Wesenberg DT, et al. Conservation of Cdc14 phosphatase specificity in plant fungal pathogens: implications for antifungal development. Sci Rep. 2020 Dec;10(1):12073.

**A**

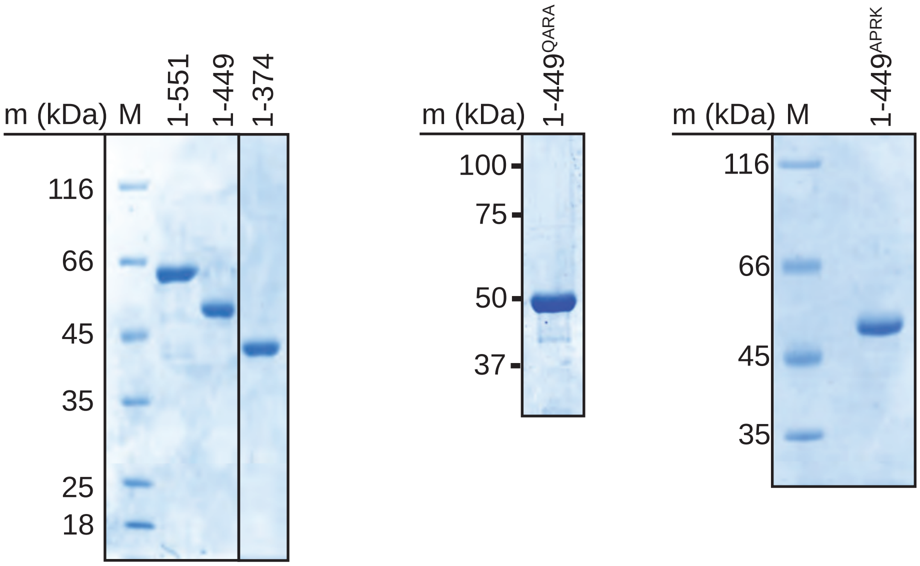


**B**

| Name | Peptide Sequence |
|---|---|
| pS | HT(pS)PIKSIG |
| pT | HT(**pT**)PIKSIG |
| pY | HT(**pY**)PIKSIG |
| +1A | HT(pS)**A**IKSIG |
| +2K | HT(pS)P**K**KSIG |
| +2R | HT(pS)P**R**KSIG |
| +3A | HT(pS)PI**A**SIG |
| +3R | HT(pS)PI**R**SIG |
| +4K | HT(pS)PIK**K**IG |
| +4R | HT(pS)PIK**R**IG |
| +2K,+3A | HT(pS)P**KA**SIG |
| +3A,+4K | HT(pS)PI**AK**IG |

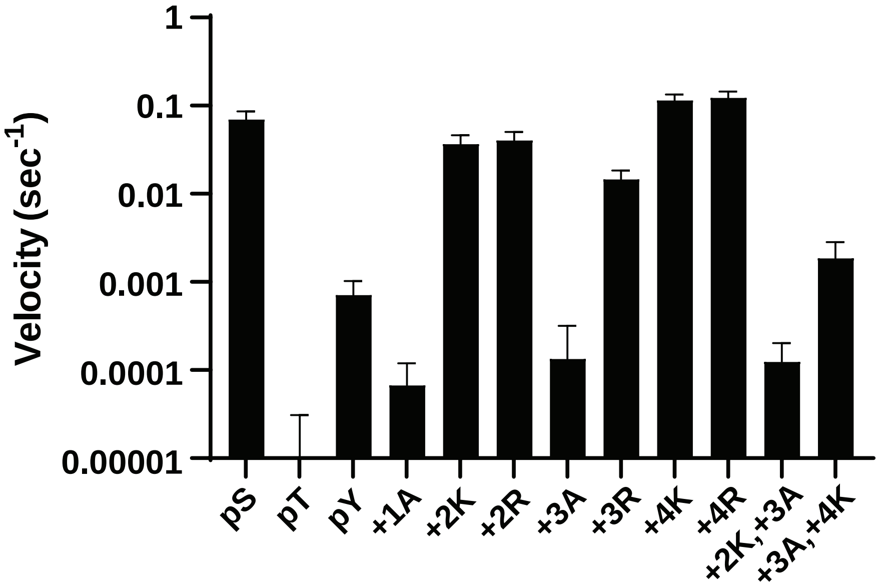


FIGURE S1

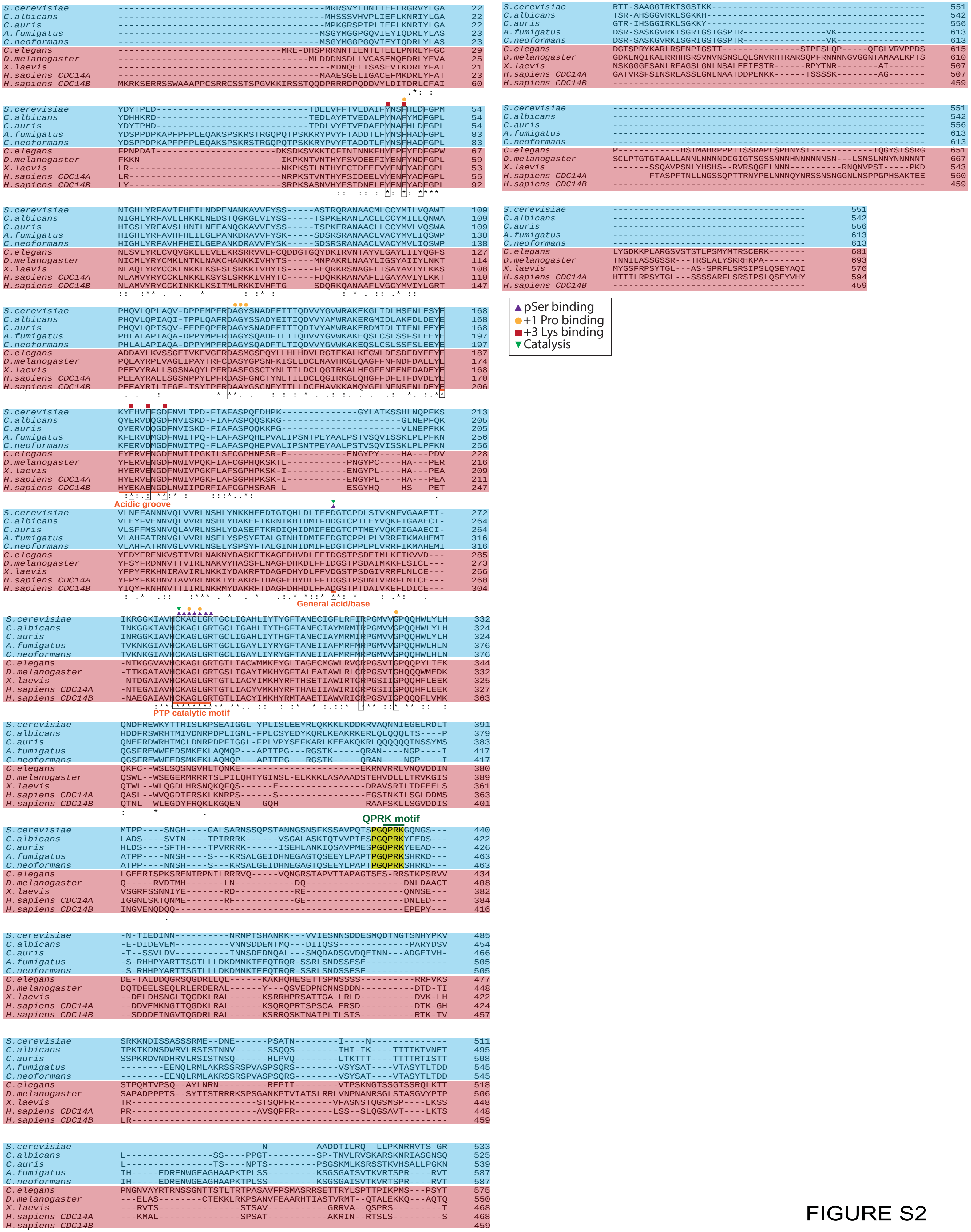
pSer binding
+1 Pro binding
+3 Lys binding
Catalysis
S.cerevisiae
C.albicans
C.auris
A.fumigatus
C.neoformans
C.elegans
D.melanogaster
X.laevis
H.sapiens CDC14A
H.sapiens CDC14B
Acidic groove
General acid/base
PTP catalytic motif
QPRK motif

FIGURE S2

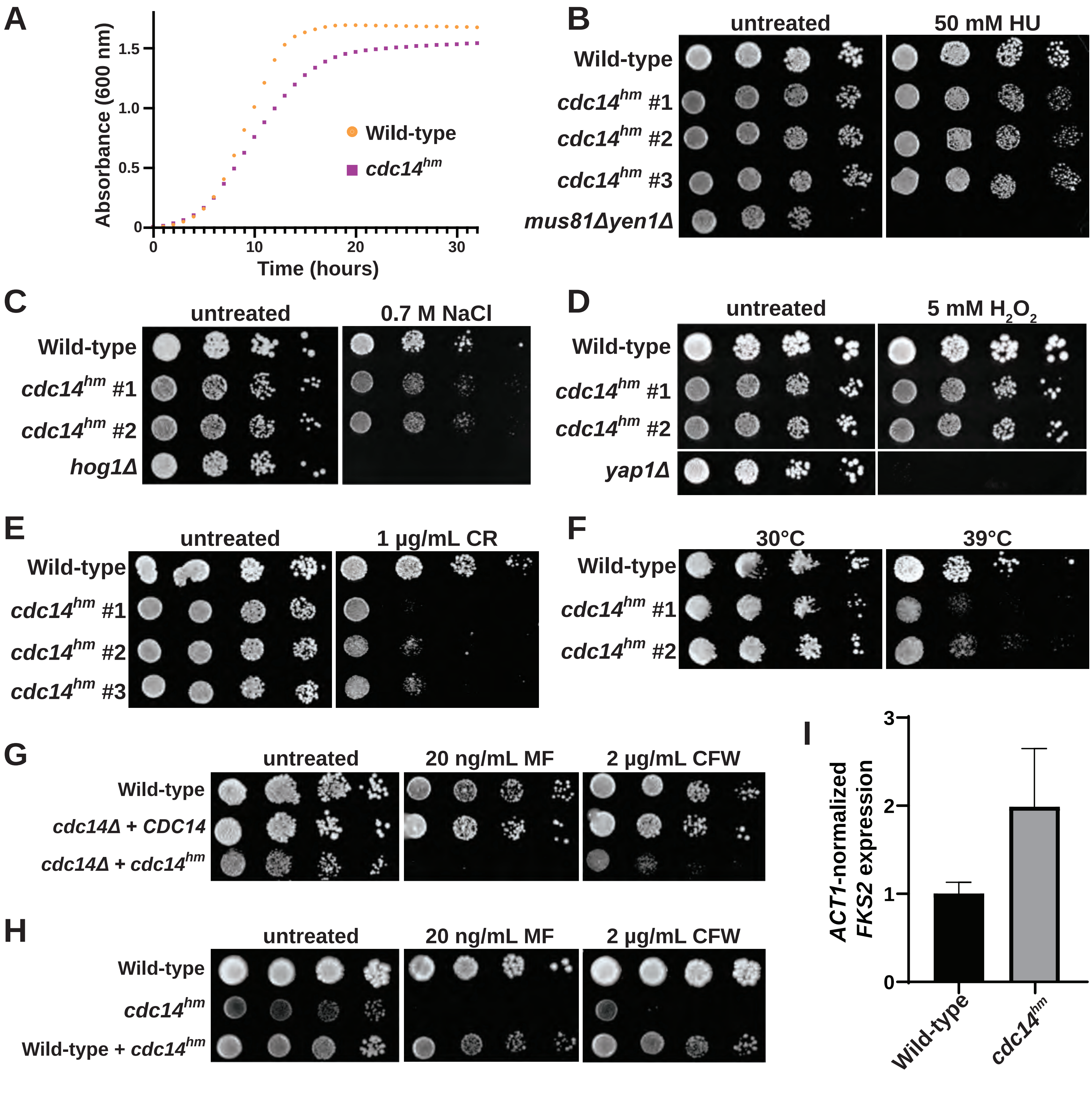
A
Absorbance (600 nm)
Time (hours)
Wild-type
cdc14hm
B
untreated
50 mM HU
Wild-type
cdc14hm #1
cdc14hm #2
cdc14hm #3
mus81Δyen1Δ
C
untreated
0.7 M NaCl
Wild-type
cdc14hm #1
cdc14hm #2
hog1Δ
D
untreated
5 mM H2O2
Wild-type
cdc14hm #1
cdc14hm #2
yap1Δ
E
untreated
1 µg/mL CR
Wild-type
cdc14hm #1
cdc14hm #2
cdc14hm #3
F
30°C
39°C
Wild-type
cdc14hm #1
cdc14hm #2
G
untreated
20 ng/mL MF
2 µg/mL CFW
Wild-type
cdc14Δ + CDC14
cdc14Δ + cdc14hm
H
untreated
20 ng/mL MF
2 µg/mL CFW
Wild-type
cdc14hm
Wild-type + cdc14hm
I
ACT1-normalized FKS2 expression
Wild-type
cdc14hm

FIGURE S3

A

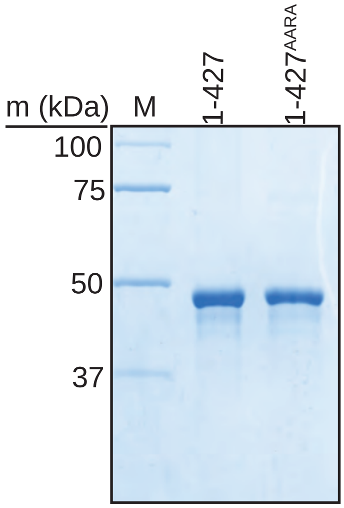


B

| Name | Peptide Sequence |
| --- | --- |
| pS | HT(pS)PIKSIG |
| pT | HT(**pT**)PIKSIG |
| pY | HT(**pY**)PIKSIG |
| +1A | HT(pS)**A**IKSIG |
| +2K | HT(pS)P**K**KSIG |
| +2R | HT(pS)P**R**KSIG |
| +3A | HT(pS)PI**A**SIG |
| +3R | HT(pS)PI**R**SIG |
| +4K | HT(pS)PIK**K**IG |
| +4R | HT(pS)PIK**R**IG |
| +2K,+3A | HT(pS)P**KA**SIG |
| +3A,+4K | HT(pS)PI**AK**IG |

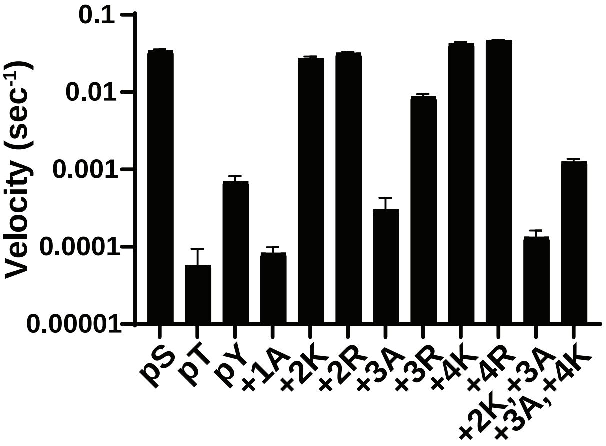


C

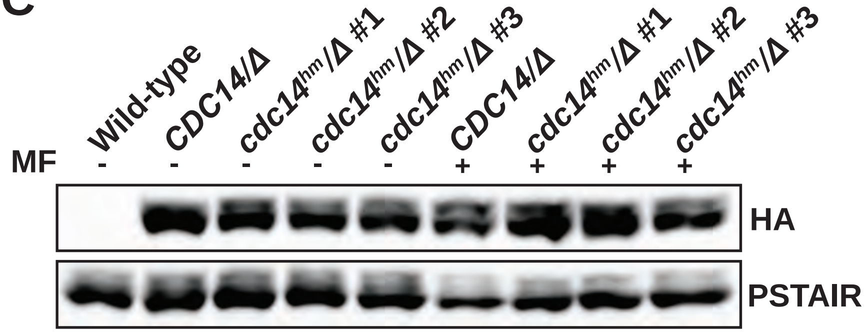


D

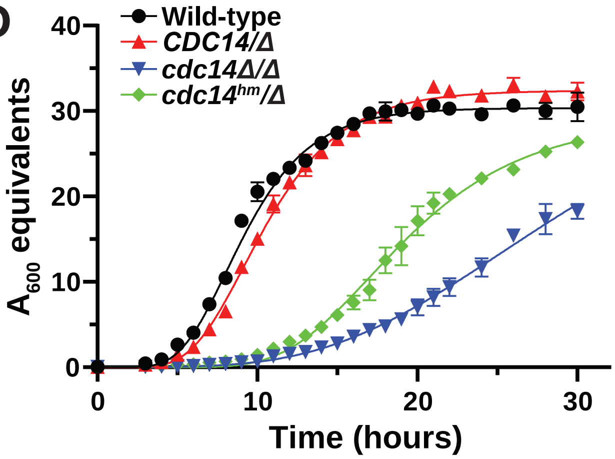


E

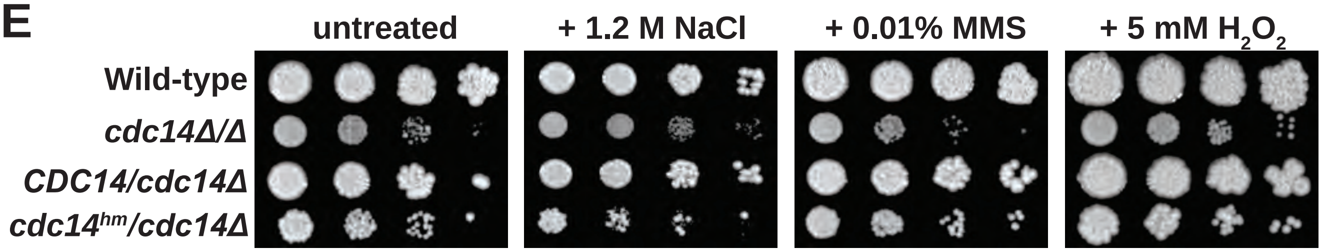


FIGURE S4

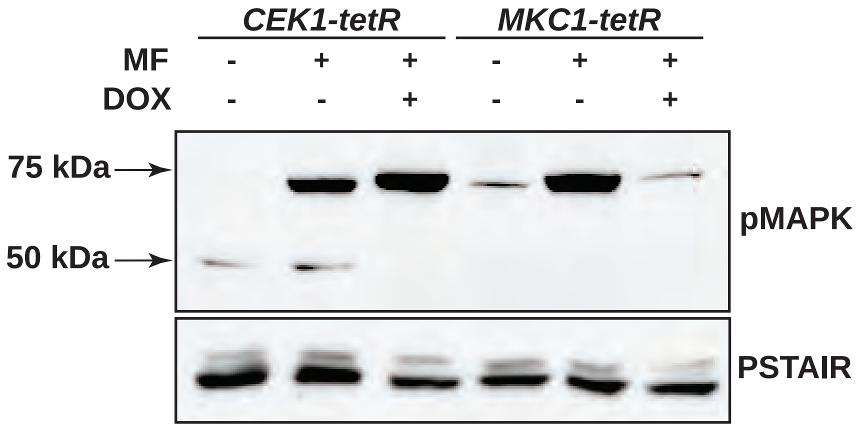


FIGURE S5

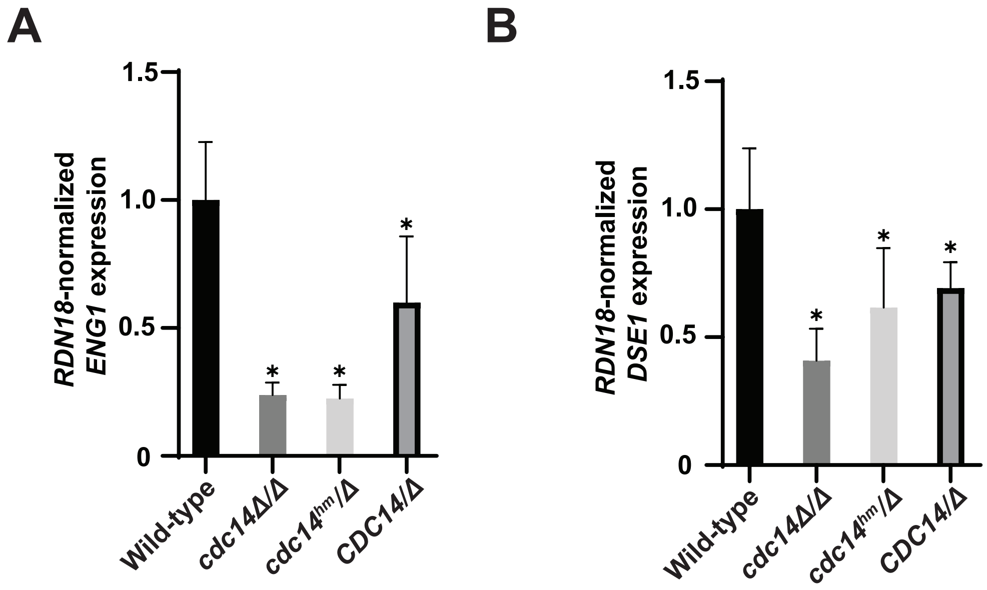
A
B
RDN18-normalized ENG1 expression
RDN18-normalized DSE1 expression
1.5
1.0
0.5
0
Wild-type
cdc14Δ/Δ
cdc14hm/Δ
CDC14/Δ
*

FIGURE S6

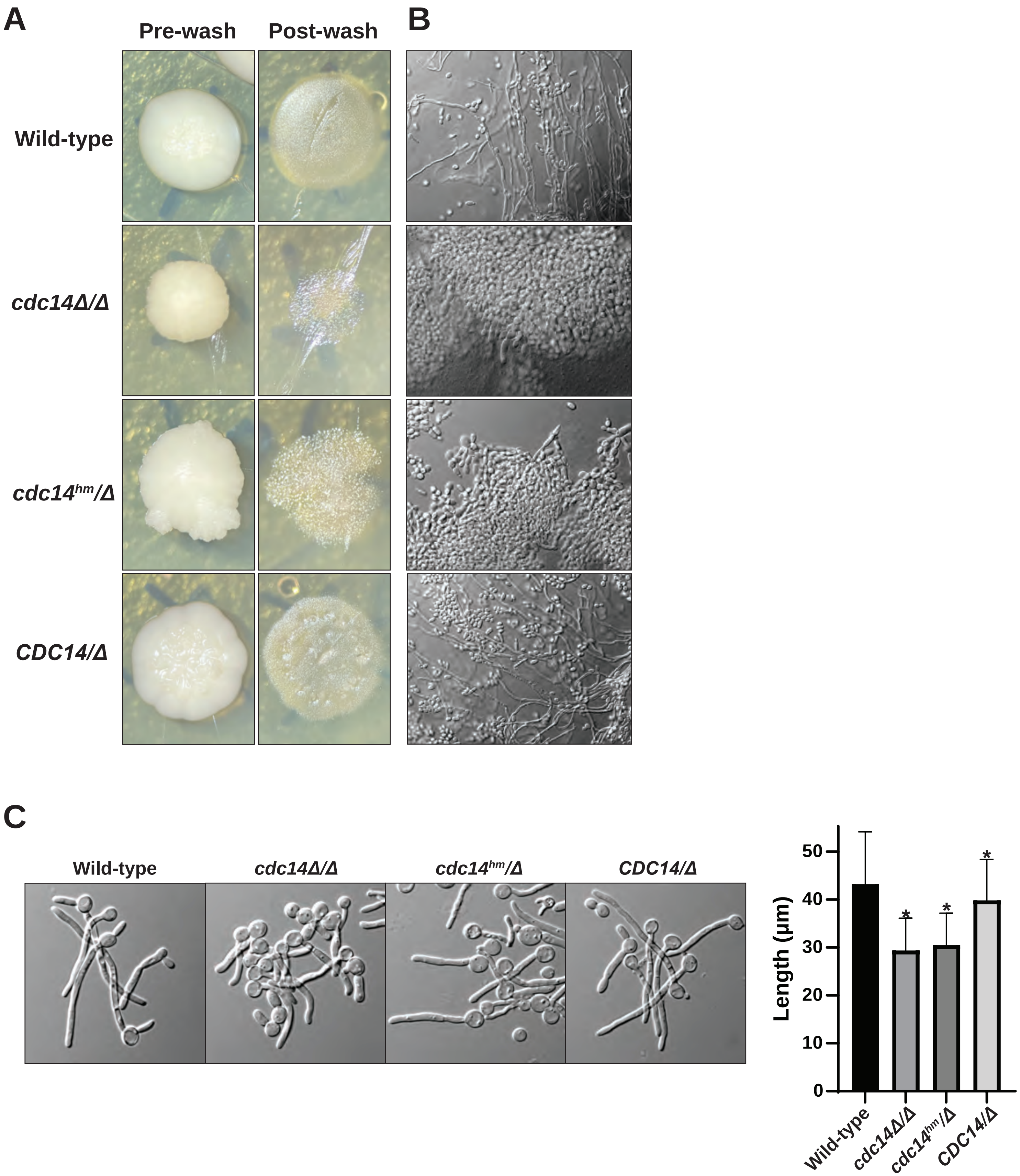
A
Pre-wash
Post-wash
B
Wild-type
cdc14Δ/Δ
cdc14hm/Δ
CDC14/Δ
C
Wild-type
cdc14Δ/Δ
cdc14hm/Δ
CDC14/Δ
Length (µm)
0
10
20
30
40
50
*
Wild-type
cdc14Δ/Δ
cdc14hm/Δ
CDC14/Δ

FIGURE S7

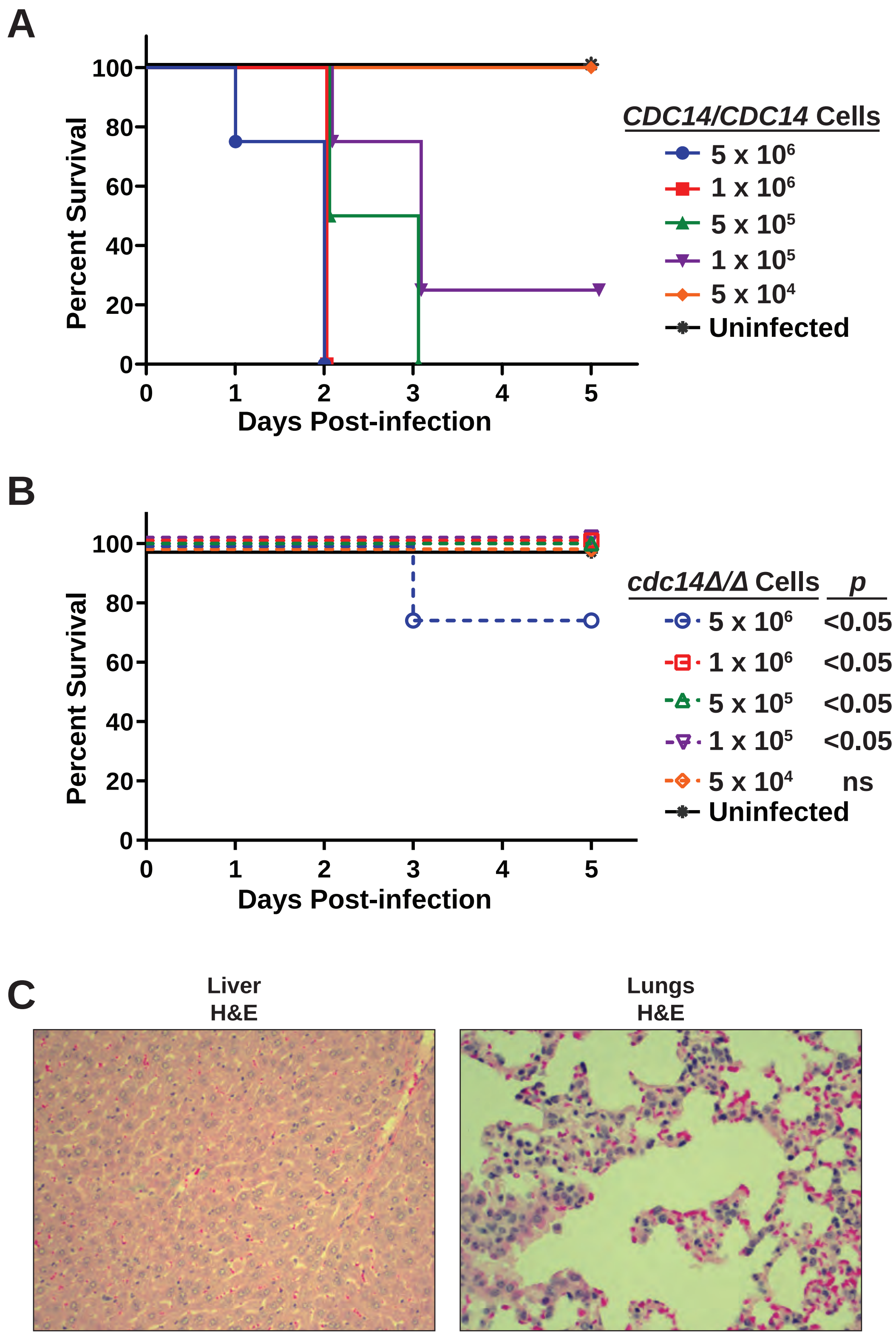
A
Percent Survival
100
80
60
40
20
0
0 1 2 3 4 5
Days Post-infection
CDC14/CDC14 Cells
5 x 10^6
1 x 10^6
5 x 10^5
1 x 10^5
5 x 10^4
Uninfected
B
Percent Survival
100
80
60
40
20
0
0 1 2 3 4 5
Days Post-infection
cdc14Δ/Δ Cells p
5 x 10^6 <0.05
1 x 10^6 <0.05
5 x 10^5 <0.05
1 x 10^5 <0.05
5 x 10^4 ns
Uninfected
C
Liver
H&E
Lungs
H&E

FIGURE S8
